# Audiovisual adaptation is expressed in spatial and decisional codes

**DOI:** 10.1101/2021.02.15.431309

**Authors:** Máté Aller, Agoston Mihalik, Uta Noppeney

## Abstract

The brain adapts dynamically to the changing sensory statistics of its environment. The neural circuitries and representations that support this cross-sensory plasticity remain unknown. We combined psychophysics and model-based representational fMRI and EEG to characterize how the adult human brain adapts to misaligned audiovisual signals. We show that audiovisual adaptation moulds regional BOLD-responses and fine-scale activity patterns in a widespread network from Heschl’s gyrus to dorsolateral prefrontal cortices. Crucially, audiovisual recalibration relies on distinct spatial and decisional codes that are expressed with opposite gradients and timecourses across the auditory processing hierarchy. Early activity patterns in auditory cortices encode sounds in a continuous space that flexibly adapts to misaligned visual inputs. Later activity patterns in frontoparietal cortices code decisional uncertainty consistent with these spatial transformations. Our findings demonstrate that regions throughout the auditory processing hierarchy multiplex spatial and decisional codes to adapt flexibly to the changing sensory statistics in the environment.

## Introduction

Throughout life the brain needs to adapt dynamically to changes in the environment and the sensorium. Changes in the sensory statistics evolve across multiple timescales ranging from milliseconds to years and bring cues from different sensory modalities into conflict. Most notably, physical growth, ageing or entering a room with reverberant acoustics can profoundly alter the sensory cues that guide the brain’s construction of spatial representations. To maintain auditory and visual spatial maps in co-registration the brain needs to constantly recalibrate the senses^1^.

Recalibrating auditory and visual spatial maps is particularly challenging, because the two sensory systems encode space not only in different reference frames (i.e., eye vs. head-centred), but also in different representational formats^2–4^. In vision, spatial location is encoded directly in the sensory epithelium and retinotopic maps of visual cortices (‘place code’)^5^. In audition, azimuth location is computed mainly from interaural time and level differences in the brain stem^2^. In primate auditory cortices, sound location is thought to be encoded by activity differences between two neuronal populations, broadly tuned to ipsi- or contralateral hemifields (i.e., ‘hemifield code’)^6–9^. Less is known about the coding principles at successive processing stages in parietal or prefrontal cortices, in which the hemifield code may be converted into a place code with narrow spatial tuning functions comparable to vision (e.g. ventral intraparietal area^10^).

Mounting behavioural research illustrates the brain’s extraordinary ability for cross-sensory plasticity. Most prominently, exposure to synchronous, yet spatially misaligned audiovisual signals biases observer’s perceived sound location towards a previously presented visual stimulus – a phenomenon coined ventriloquist aftereffect^11–14,3,15–19^. This cross-sensory adaptation to intersensory disparities emerges at multiple timescales ranging from milliseconds^13,20,18^ to minutes^3,16,17,19^ or even days^21^.

Yet, the underlying neural circuitries, mechanisms and representations that support cross-sensory plasticity remain unknown. The frequency selectivity of audiovisual recalibration has initially pointed towards early stages in tonotopically organized auditory cortices (see^16^ but^14^). By contrast, the involvement of hybrid spatial reference frames that are neither eye- nor head-centred suggested a pivotal role for parietal cortices or inferior colliculus^3,22^.

Importantly, recalibration may arise at multiple levels affecting spatial and choice-related computations. While the former alters neural encoding of sound location irrespective of task, the latter affects the read out of decisional choices from those neural representations. The dissociation of the two is challenging for behavioural research that is forced to estimate recalibration from behavioural responses. Likewise, previous neuroimaging studies were not able to disentangle the two, because they employed a spatial localization task that maps each sound location onto one particular response choice, thereby conflating spatial and choice-related processes^13,19,20^. To dissociate changes in spatial and decisional representations, we need neuroimaging experiments that map different sound locations onto identical decisional choices^17^.

This study investigates how changes in audiovisual statistics mould auditory spatial and decisional coding. Participants performed a spatial classification task on auditory stimuli before and after audiovisual recalibration. Combining model-based analyses of regional BOLD-responses and fine-scaled EEG and fMRI activity patterns, we temporally and spatially resolved how the brain flexibly adapts spatial and decisional coding to changes in the sensory statistics.

## Results

In each participant, we performed psychophysics, functional magnetic resonance imaging (fMRI) and electroencephalography (EEG) experiments on 13 days (Figure 1A). Each experiment included unisensory auditory pre-adaptation, left (i.e., VA) and right (i.e., AV) audiovisual adaptation, and auditory postVA/postAV-adaptation phases. During pre- and post-adaptation phases, participants were presented with auditory stimuli at 7 spatial locations (±12°, ±5°, ±2° and 0°) along the azimuth. They performed a left-right spatial classification task only on 22% ‘response trials’ that were randomly interspersed in ‘non-response trials’ (Figure 1B). During the audiovisual adaptation phases, participants were presented with a sound in synchrony with a visual stimulus that was displaced in separate sessions by 15° to the left or right of the sound location. To focus on implicit perceptual recalibration, observers performed a non-spatial task during the adaptation phases: they detected small changes in contrast of the visual stimulus that occurred on 10% ‘response trials’ (see Figure 1B). Direct comparison of postVA and postAV adaptation trials enables the assessment of recalibration unconfounded by time and learning effects.

**Figure 1.**
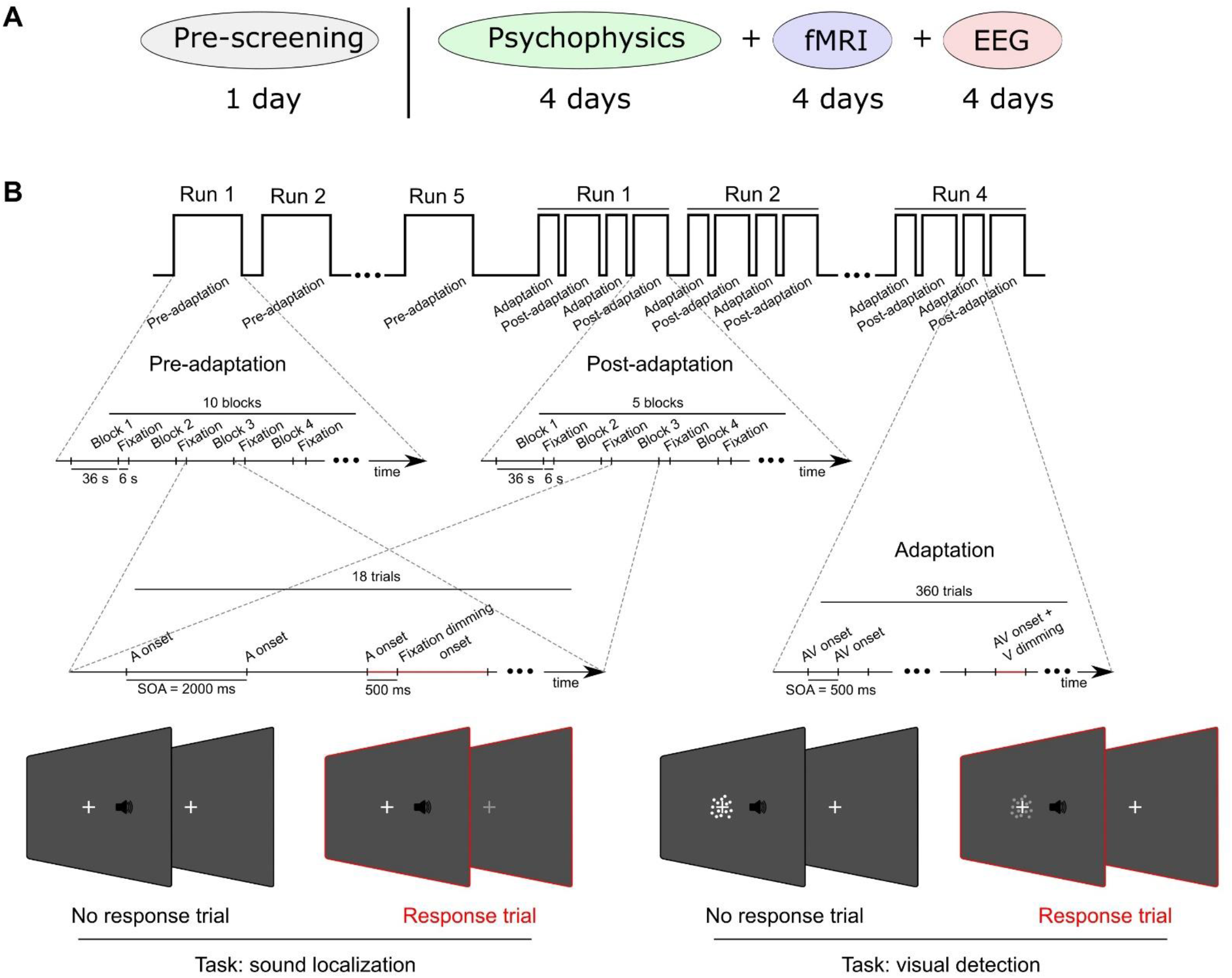
Study and experimental design. **(A)** The study included one day of pre-screening, four days of psychophysics testing, four days of fMRI and four days of EEG. **(B)** Each day included (i) auditory pre-adaptation, (ii) left (i.e., VA) or right (AV) audiovisual adaptation and (iii) auditory post-adaptation phases. In the pre-adaptation and post-adaptation phases, observers were presented with unisensory auditory stimuli sampled from seven (±12°, ±5°, ±2° and 0° visual angle) locations along the azimuth. They performed a spatial (left vs. right) classification task on 22% of the trials (‘response trial’) that were indicated by a brief dimming of the fixation cross 500 ms after sound onset. In the audiovisual adaptation phases, observers were presented with audiovisual stimuli with a spatial disparity of ±15°: the visual signal was sampled from three locations along the horizontal plane (i.e., −5°, 0°, 5°) and the visual signal was spatially shifted by 15° either to the left (i.e., VA-adaptation) or to the right (i.e., AV-adaptation) with respect to the auditory stimulus. Observers were engaged in a non-spatial visual detection task on 10% of the trials (‘response-trial’) indicated by the lower contrast of the visual signals. Each day started with the pre-adaptation phase. Next, VA (or AV)-adaptation phases alternated with postVA (or postAV)-adaptation phases. This figure shows the experimental details for the fMRI experiments, which were slightly modified for EEG and psychophysics experiments (for details see Experimental design and procedure section).

### Behavioural results

Throughout all experiments, observers were attentive and followed task instructions as indicated by a sensitivity index (d’) of > 4 across all experimental phases (Table S1). Observers responded to > 88% of ‘response trials’ (i.e., ‘hits’) both in the pre-/post-adaptation (i.e., 22% response trials) and the adaptation phases (i.e., 10% response trials, Table S1). They made only < 0.47% false alarms (i.e., responses to non-response trials).

Likewise, eye movement analyses during the psychophysics experiment showed that fixation was well maintained on 94.02% ± 1.00% (mean ± SEM) of the trials throughout the entire experiment with no significant differences in eye movement indices between postVA- and postAV-adaptation phases (i.e., % saccades, % eye blinks, and post-stimulus mean horizontal fixation position, see Supplementary Results). Potential recalibration effects are therefore not confounded by eye movements.

To assess whether exposure to misaligned visual signals recalibrates observers’ sound representations, we fitted cumulative Gaussians as psychometric functions to the percentage ‘perceived right’ responses in the auditory post-adaptation phases (Figure 2A). Consistent with previous research^15–17^ observers’ perceived sound locations shifted towards the displaced visual stimulus that was presented in the prior adaptation phases. Bayesian model comparison confirmed that the recalibration model that allows for shifts in the point of subjective equality (PSE) across pre-, postVA-, and postAV-adaptation phases was exceedingly more likely than the static model in which the PSE values were constrained to be equal (protected exceedance probabilities > 0.88 in each of the psychophysics, fMRI, and EEG experiments; Table S2). Moreover, paired t-tests of the PSE values (from the recalibration model) between postVA- and postAV-adaptation phases revealed significant differences in PSE between postVA- and postAV-adaptation phases (psychophysics: t(14) = 11.4, p < 0.0001; fMRI: t(4) = 11.3, p < 0.001; EEG: t(4) = 10.4, p < 0.001) (Figure 2B). Collectively, our behavioural results confirmed that prior exposure to disparate audiovisual signals recalibrate observers’ spatial classification responses to sounds.

**Figure 2.**
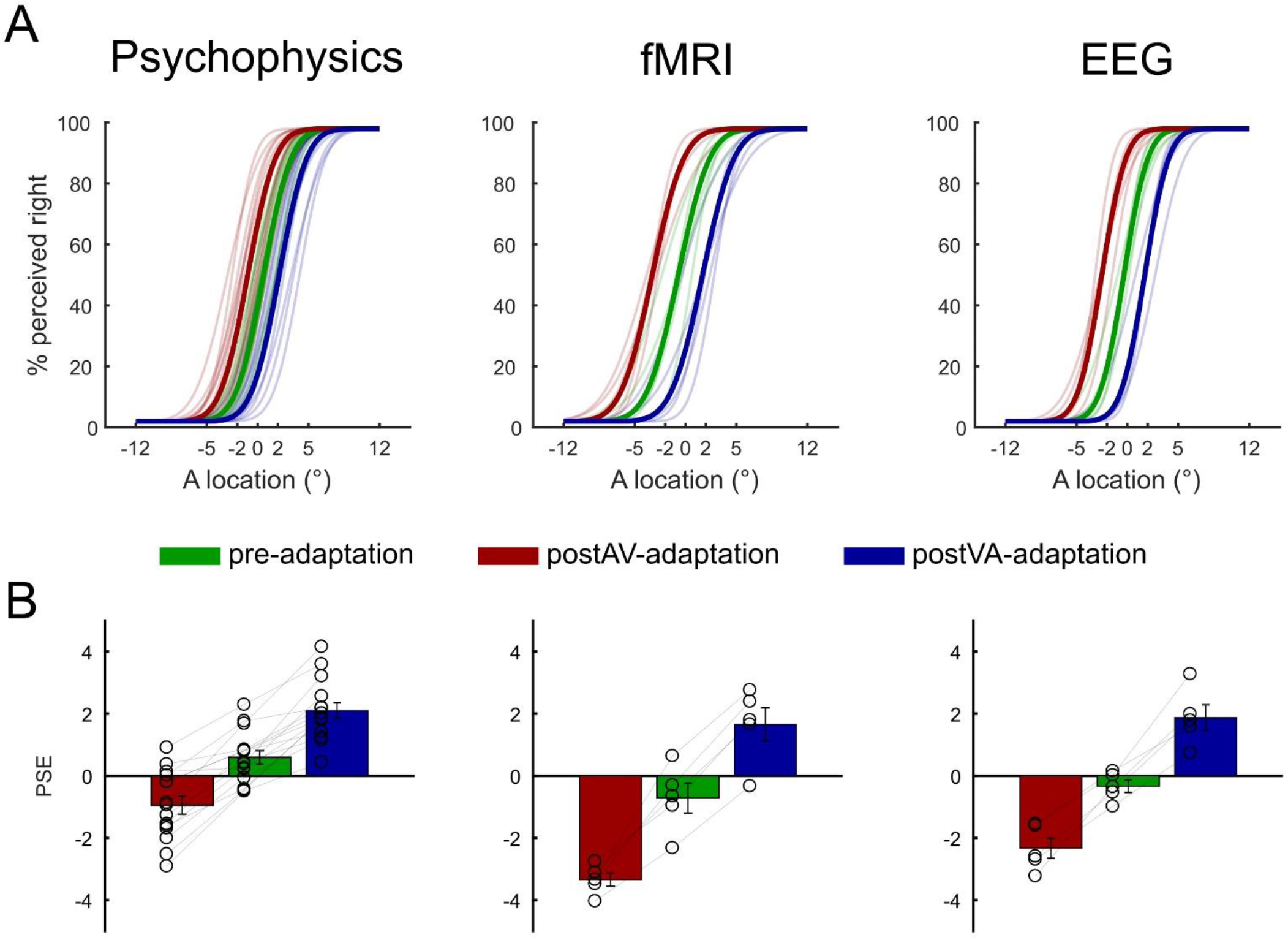
Behavioural results. **(A)** Psychometric functions fitted to the behavioural data from the psychophysics, fMRI, and EEG experiments for pre- (green), postAV- (red), and postVA-adaptation (blue) phases (thin lines for subject-specific psychometric functions, thick line for across-subjects’ mean). **(B)** Across-subjects’ mean (± SEM) of the PSE for pre-adaptation (green), postVA- (red) and postAV-adaptation (blue) phases. The subject-specific PSEs are overlaid as circles. The PSE was shifted towards the left for postAV- (red) relative to postVA-adaptation (blue) phases in every single participant. PSE: point of subjective equality, SEM: standard error of the mean.

### fMRI results

#### Spatial encoding and recalibration indices

Using multivariate decoding we identified brain areas that (i) encoded sound location and (ii) recalibrated this encoded sound location. Guided by previous research^7,19,23–26^ we focused on five regions of interest (ROI): Heschl’s gyrus (HG), higher auditory cortex (hA, mainly planum temporale), intraparietal sulcus (IPS), inferior parietal lobule (IPL) and frontal eye field (FEF). In each of those ROIs we trained a linear support vector regression model (SVR, LIBSVM^27^) in a 4-fold cross-validation scheme to learn the mapping from BOLD-response patterns of the pre-adaptation phase to external auditory locations. We used this learnt mapping to decode the sound location from the activation patterns of the pre-, postVA-, and postAV-adaptation phases. We observed significantly better than chance decoding accuracies (i.e., ‘spatial encoding index’, the Fisher z-transformed Pearson correlation coefficient between the true and the decoded sound locations) across all ROIs along the auditory spatial processing hierarchy with a maximal decoding accuracy in planum temporale (right-tailed p-values for the second level bootstrap-based one sample t-tests: HG: p = 0.028; hA: p = 0.012; IPS: p = 0.005; IPL: p = 0.010; FEF: p = 0.019, see Figure 3A).

**Figure 3.**
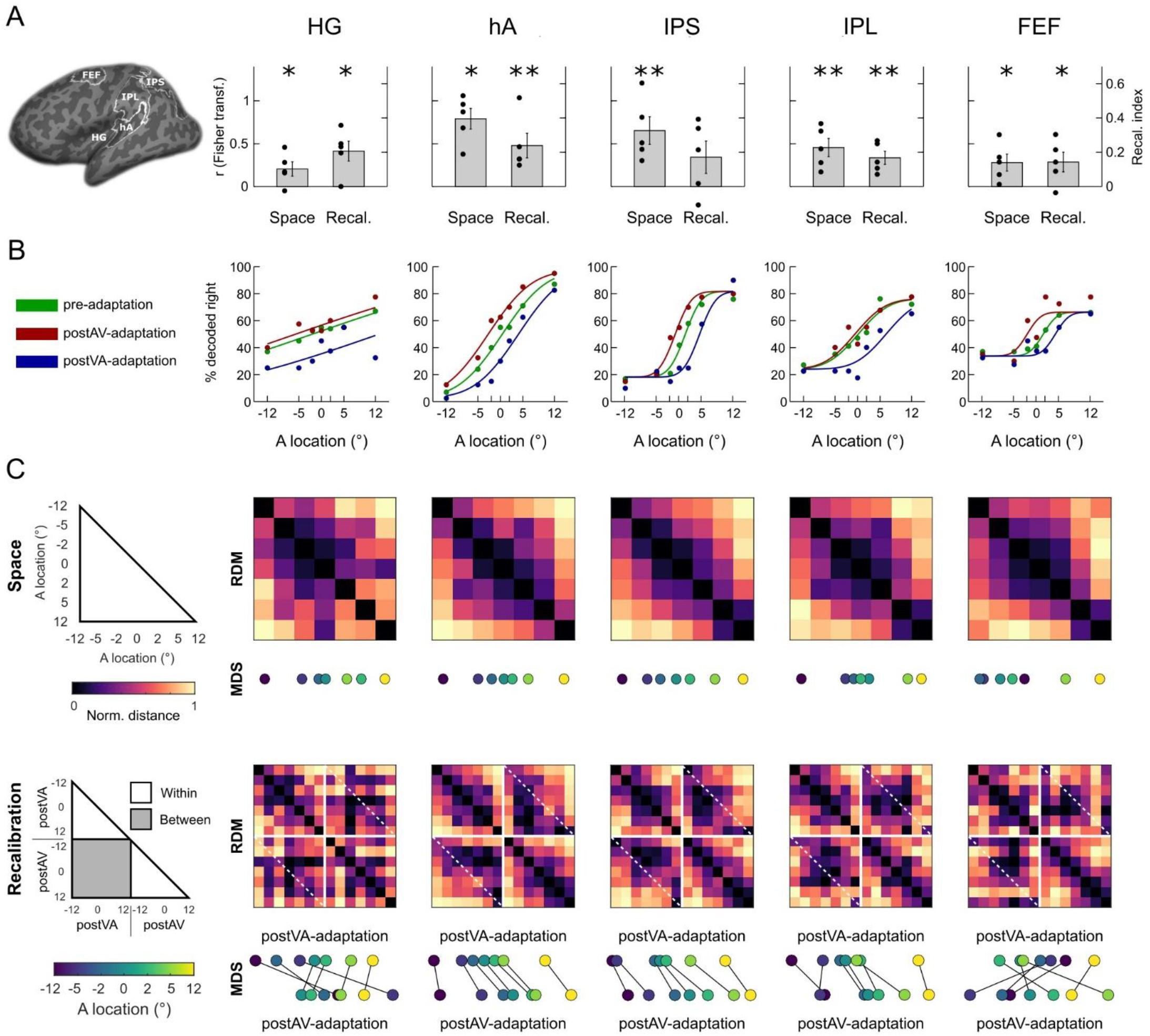
fMRI multivariate decoding and representational dissimilarity matrices. **(A)** Across-subjects’ mean (± SEM) of the spatial encoding (left bars, Fisher z-transformed correlation coefficient) and recalibration (right bars, difference in fraction ‘decoded right’ for postAV-adaptation vs. postVA-adaptation) indices in the ROIs as indicated in the figure. *: p ≦ 0.05; **: p ≦ 0.01. The subject-specific values are overlaid as black dots. **(B)** Neurometric function fitted to the fMRI decoded sound locations across ROIs for pre-, postVA-, and postAV-adaptation phases. **(C)** RDMs (across-subjects’ mean) showing the Mahalanobis distances for the fMRI decoded sound locations across ROIs. Top row: 7 × 7 RDM for pre-adaptation phase. Bottom row: 14 × 14 matrix for postVA- and postAV-adaptation. The drawings on the left indicate the organization of the RDMs of spatial locations for pre- (top) as well as for postAV- and postVA-adaptation phases (bottom). Using MDS we projected each RDM onto a single dimension (i.e., representing space along the azimuth). For illustration purposes, we vertically offset the MDS projections for postVA- and postAV-adaptation phases and connected the projections corresponding to the same physical stimulus locations. The MDS results show that the projected locations of the postVA-adaptation phase are shifted towards the left in relation to those of the postAV-adaptation phase particularly in hA and IPS, while a more complex relationship arises in IPL and FEF. HG: Heschl’s gyrus; hA: higher auditory cortex; IPL: inferior parietal lobule; IPS: intraparietal sulcus; FEF: frontal eye-field; RDM: representational dissimilarity matrix; MDS: multidimensional scaling.

To assess whether these encoded sound locations adapt to displaced visual stimuli, we computed the recalibration index (RI) as the difference between the fraction of ‘decoded right’ in the postAV-adaptation phase minus the postVA-adaptation phase. If the decoded sound location shifts towards the disparate visual signal during the AV- and VA-adaptation phase, we would expect a positive recalibration index. Consistent with this conjecture we observed a RI that was significantly greater than zero in nearly all ROIs (right-tailed p-value of second level bootstrap-based one sample t-test, HG: p = 0.025, hA: p = 0.002, IPL: p = 0.010, FEF: p = 0.037, and a trend in IPS: p = 0.086, see Figure 3A).

These results provide initial evidence that regions along the dorsal auditory processing hierarchy show an overall effect of recalibration averaged across all sound locations.

#### Neurometric functions

Similar to our behavioural analysis, we fitted cumulative Gaussians as neurometric functions to the percentage ‘decoded right’ separately for the pre-, postVA-, and postAV-adaptation phases at the group level (see Figure 3B). Consistent with our behavioural findings the neurometric functions shifted rightwards after VA-adaptation and leftwards after AV-adaptation. Bayesian model comparison^28^ provided decisive evidence^29^ for the recalibration model that allows for changes in PSE values between pre, postVA-, and postAV-adaptation phases relative to a static model in which the PSE values were constrained to be equal (Akaike criterion > 4.6 in all ROIs; HG: 12.9, hA: 20.2, IPS: 17.0, IPL: 10.3, FEF: 5.7; Table S3).

#### Representational similarity analysis

Our linear decoding results indicate that auditory representations throughout the processing stream are altered by prior AV recalibration. Critically, our results so far are agnostic about the coding principles underlying this pervasive cross-sensory plasticity. Next, we therefore characterized the geometry of the neural representations across the seven sound locations using representational similarity analysis (RSA^30^). For visualization, we projected the group level representational dissimilarity matrices (RDMs, averaged across participants) with non-classical multidimensional scaling (MDS) onto a single dimension to incorporate the spatial organization along the azimuth (i.e., as physical space, see Figure 3C). Particularly, higher auditory cortex (hA, including planum temporale) and IPS encoded the seven sound locations largely consistent with the physical distances of the sound locations. For instance, in both hA and IPS, the MDS distance is greater for the sound locations 12° and 5° than for the locations 5° and 0°. By contrast, the representational geometry in IPL and FEF does not fully correspond to the physical sound distances (Figure 3C bottom row). Across all regions, the MDS shows spatial shifts to the left (resp. right) after AV- (resp. VA)-adaptation. Again, the similarity between neural representations in hA and IPS obeyed the physical order of the sound locations, while this is less clear for IPL or FEF. For instance, in FEF the neural representations for the ‘-12° sound’ is shifted towards the centre (see Figure 3C, FEF in bottom row).

#### Model-based fMRI analysis: dissociating perceptual and decisional codes

MDS revealed subtle differences in representational geometry across regions. These may reflect noise or arise from mixing of multiple representational components. Most notably, recalibration may affect spatial and decisional (i.e., choice-related) representations. To arbitrate between these hypotheses, we compared a spatial, a decisional or a combined spatial + decisional model as explanations for the regional mean BOLD-response and/or the fine-scale activation patterns.

To model spatial/perceptual representations we used the hemifield model (see Figure 4A) that encodes sound location in the relative activity of two subpopulations of neurons, each broadly tuned to the ipsi- or contra-lateral hemifield^6,31^. Because the hemifield and place code models make near-indistinguishable predictions for the pattern similarity structure over the relatively central locations used in the current study, we applied the hemifield model as a generic spatial model to all regions (for details, see Supplementary Methods). The decisional model (see Figure 4A) codes observers’ decisional uncertainty as a non-linear function of the distance between the spatial estimate from the left/right classification boundary^32^ (see Methods for further details).

**Figure 4.**
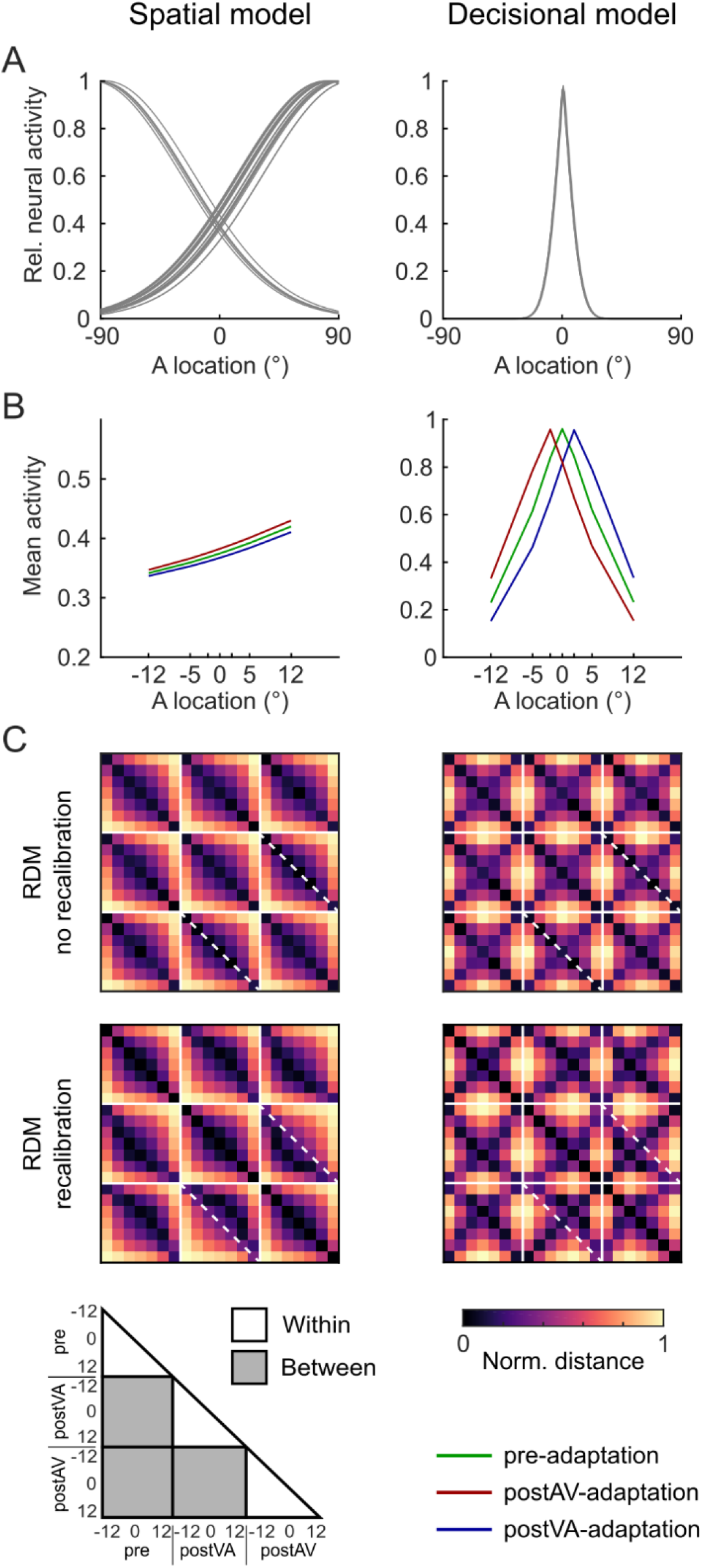
Spatial and decisional models. **(A)** Neural models: The spatial (hemifield) model encodes spatial location in the relative activity of two subpopulations of neurons each broadly tuned either to the ipsi- or contralateral hemifield. The ratio of the ipsi- and contralaterally tuned neurons was set to 30%/70% consistent with prior research^8^. The decisional model encodes observers’ decisional uncertainty as a non-linear function of the distance between the spatial estimates and the spatial classification boundary. **(B)** Predicted mean BOLD-response as a function of sound location along the azimuth in a left hemisphere region for pre- (green), postVA- (blue) and postAV-adaptation (red). The spatial model predicts a BOLD-response that increases linearly for sound locations along the azimuth. The decisional model predicts BOLD-response that decays with the distance from the decision boundary in an inverted U-shaped function. Further, this inverted U-shaped function shifts along the azimuth when spatial estimates are recalibrated. The model predictions were obtained by averaging the simulated neural activities across 360 neurons in a left hemisphere region. **(C)** Predicted representational dissimilarity matrices (RDM) based on the individual model neural activity profiles across spatial locations (−12° to 12°) and experimental phases (pre, postVA and postAV). We simulated RDMs from the spatial (left) and the decisional (right) model for (i) top row: no recalibration, i.e., without a representational shift and (ii) bottom row: with recalibration, i.e., with a representational shift. Solid white lines delineate the sub-RDM matrices that show the representational dissimilarities for different stimulus locations within and between different experimental phases. Diagonal dashed white lines highlight the RDM dissimilarity values for two identical physical locations of the postAV- and the postVA-adaptation phases. Comparing the RDMs with and without recalibration along those dashed white lines shows how the shift in spatial representations towards the previously presented visual stimulus alters the representational dissimilarity of corresponding stimulus locations in postAV- and postVA-adaptation phases, while the off-diagonals show the dissimilarity values for neighbouring spatial locations.

Because a region may combine spatial and decisional coding, we first assessed whether (i) the spatial model (S), (ii) the decisional model (D) or (iii) the combined spatial + decisional model (S+D) were the best explanations for our data. In a second step, we used the combined spatial + decisional model to assess the contributions of the spatial and decisional components to recalibration by comparing: (i) spatial + decisional with no recalibration in either model component (S+D), (ii) spatial with recalibration + decisional (S_R_+D), (iii) spatial + decisional with recalibration (S+D_R_) and (iv) spatial with recalibration + decisional with recalibration (S_R_+D_R_). Each model component accounted for recalibration by shifting the ‘encoded sound locations’ by a constant ±2.3° (i.e., across-subjects’ mean in behavioural PSE shift) to the right (postVA-adaptation) vs. left (postAV-adaptation). As shown in Figure 4, the spatial and decisional models make distinct predictions for the regional mean BOLD-response (Figure 4B) and the similarity structure of activity patterns over sound locations (Figure 4C).

##### Regional mean BOLD-response – Linear mixed effects modelling

The spatial model predicts that the regional mean BOLD-response of - for instance - a left hemisphere region increases for stimuli towards the right hemifield. Likewise, its response to all sounds irrespective of location should be greater after right recalibration. By contrast, the decisional model predicts a mean BOLD-response that peaks at the decisional boundary and tapers off with greater distance from the boundary. As a result of recalibration, this peak shifts towards the left or right, because the brain’s spatial estimates have been recalibrated leading to a change in the relative distance between the spatial estimates and the decision boundary (see Figure 4B). For instance, the encoded spatial estimate for a sound stimulus at physical location of −5° may be shifted towards the centre after right (i.e., AV) recalibration and thereby be associated with a greater decisional uncertainty.

In hA the mean BOLD-response increased progressively for stimuli along the azimuth as expected under the spatial model (Figure 5A). By contrast, in IPS, IPL and FEF the BOLD-response peaked for physical 0° sound location in the pre-adaptation phase, left sound locations (i.e., < 0°) in the postAV-adaptation and right sound locations (i.e., > 0°) in the postVA-adaptation phase as expected under the decisional model. This visual impression was confirmed by Bayesian model comparison of the three (i.e., spatial, decisional, spatial + decisional) linear mixed effects (LME) models. Each LME model used the simulated activity of the spatial and/or decisional models as fixed effects predictors (in addition to a constant regressor) for the 7 (sound locations) × 3 (pre-, postAV-, and postVA-adaptation) conditions. In HG the most parsimonious baseline model outperformed the other models suggesting that the regional BOLD-response in HG did not provide reliable information about sound location. The Bayes factors provided strong evidence for the spatial model in hA and for the decisional model in IPS (see Table S4). Turning to the recalibration results we observed a small trend towards spatial coding in hA (i.e., S_R_+D and S_R_+D_R_ > S+D and S+D_R_), but substantial evidence towards decisional coding in IPS, IPL and FEF (i.e., S_R_+D_R_ and S+D_R_ > S+D and S_R_+D, Figure 5B bottom row, Table S4). This suggests that the mean regional BOLD-response follows the predictions of the hemifield model only in hA. By contrast in IPS, IPL and FEF the regional mean BOLD-response mainly reflects choice-related activity invoked by mapping the recalibrated spatial estimates onto observers’ decisional boundary.

**Figure 5.**
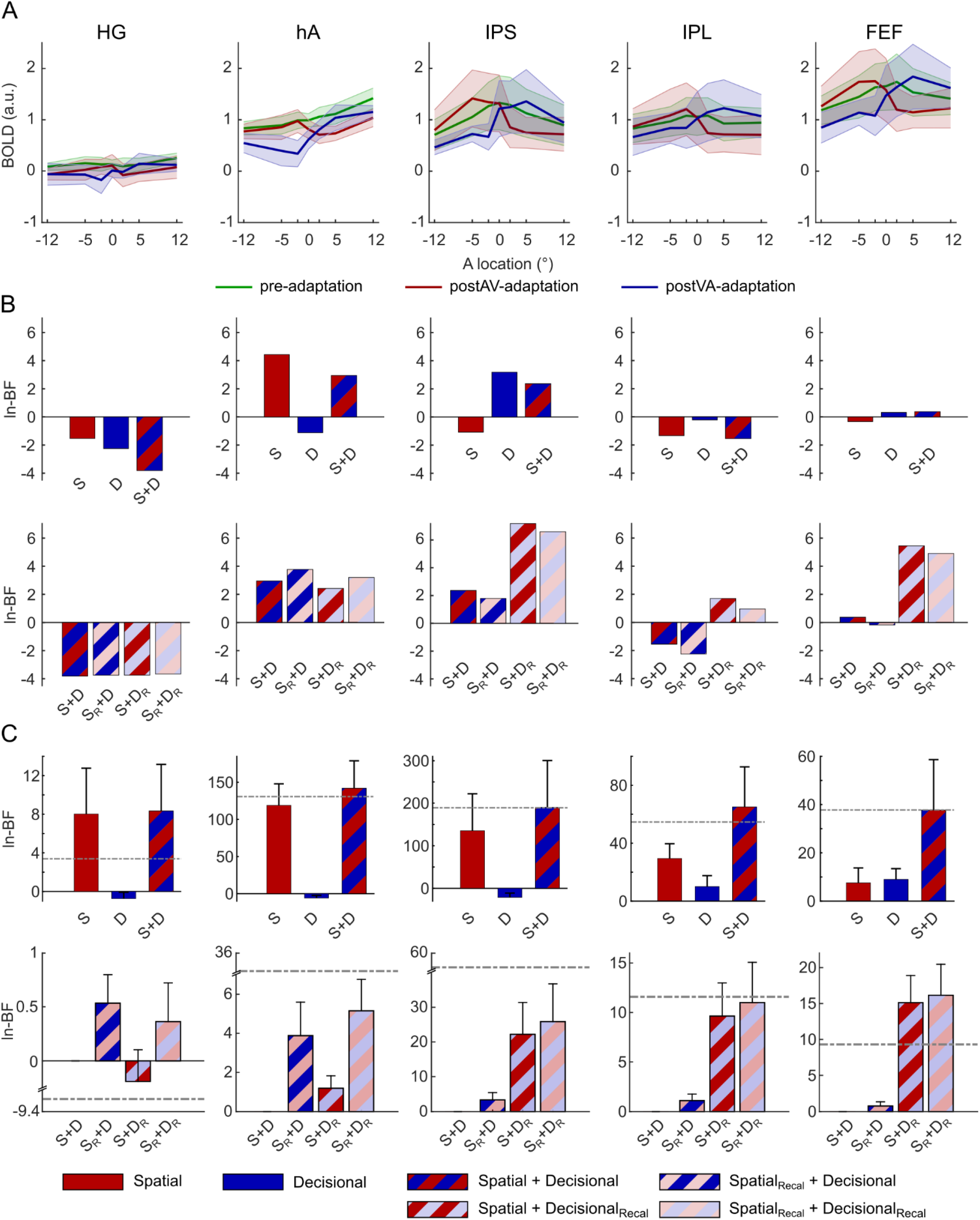
fMRI results: regional BOLD-response and pattern component modelling (PCM). **(A)**Across-subjects’ mean positive BOLD-responses shown as a function of spatial location in pre-, postVA-, and postAV-adaptation phases across ROIs. The shaded areas indicate SEM. **(B)** Results of the linear mixed effects analysis of regional mean BOLD-response across ROIs: ln-Bayes factors (ln-BF) for a specific target model as indicated by the capital letter relative to the null model, which includes only the constant term (see Supplementary methods). Top row: The spatial, decisional, and spatial + decisional models without recalibration. Bottom row: The models factorially manipulate whether the spatial and/or decisional component include recalibration. **(C)** PCM results - Spatial and/or decisional models as predictors for fine grained BOLD-response patterns across ROIs. Across-subjects mean ln-BFs (±SEM) for each target model as indicated by the capital letter. (i) Top row: The spatial, decisional, and spatial + decisional models without recalibration relative to a null model that allows for no similarity between activity patterns. (ii) Bottom row: The models factorially manipulate whether the spatial and/or decisional component account for recalibration. Ln-Bayes factors are relative to the spatial + decisional model (without recalibration) as the null model. Dash-dotted grey lines indicate the relative ln-BF for the fully flexible models as baseline estimated separately for the top and bottom row model comparisons (for details, see Methods). S = spatial model without recalibration, D = decisional model without recalibration, S_R_ = spatial model with recalibration, D_R_ = decisional model with recalibration^33^. Ln-Bayes factors of > 1.1 are considered substantial evidence and > 3 as strong evidence for one model over the other^29^.

##### Fine-scale activation pattern – Pattern component modelling

Using pattern component modelling (PCM^33^) we investigated whether spatial and decisional codes contributed to the fine-scale voxel-level activation patterns over the 7 (sound locations) × 3 (pre-, postAV-, and postVA-adaptation) conditions. Consistent with our analysis of the regional mean BOLD-response we fitted and compared a baseline model that assumes independence of activation patterns across conditions with the spatial, decisional, and combined spatial + decisional PCM models. In a second step we used the combined spatial + decisional model to assess whether recalibration was expressed in spatial, decisional, or spatial + decisional codes (see Figure 5C). We fitted all models (e.g., spatial and/or decisional etc.) and a fully flexible model using a leave-one-subject-out cross-validation scheme. The fully flexible model accommodates any possible covariance structure of activity profiles and enables us to assess whether a particular target model accounts for the key representational structure in the observations given the measurement noise and inter-subject variability (see Figure 5C).

As shown in Figure 5C top row, the ln-Bayes factors provided strong evidence for the spatial relative to the decisional model in HG and hA (see Table S5 for exact values). In HG, the spatial component alone was sufficient to reach the threshold set by the fully flexible model. In all other regions, i.e., hA, IPS, IPL and FEF, the combined spatial + decisional model outperformed the single component models suggesting that in those regions spatial and decisional representations together contribute to adaptive coding. Moreover, the decisional component became progressively more dominant along the dorsal processing hierarchy. In FEF the decisional model even outperformed the spatial model.

The recalibration results further emphasized these opposite gradients for spatial and decisional coding along the auditory processing stream. As shown in Figure 5C bottom row, in HG and hA the models expressing recalibration in spatial coding (S_R_+D and S_R_+D_R_) outperformed those without recalibration in spatial coding (S+D and S+D_R_). The evidence was substantial in hA, but not conclusive in HG. Conversely in IPS, IPL and FEF we observed strong evidence for the models that expressed recalibration in the decisional code (S+D_R_ and S_R_+D_R_) relative to those that did not (S+D and S_R_+D, see Table S5 for exact values).

In summary, our advanced model-based representational analyses demonstrate that audiovisual adaptation relies on plastic changes in spatial and decisional representations. While all regions (apart from HG) multiplexed spatial and decisional coding, their contributions arose with opposite gradients along the dorsal auditory hierarchy. The spatial model provided a better explanation for activations in HG and hA, the decisional model dominated in IPS, IPL and FEF. Crucially, only characterizing the representational geometry of the neural representations enabled us to reveal this double dissociation in spatial and decisional codes. This highlights the critical importance to move beyond simple linear decoding analyses.

### EEG results

Using EEG, we characterized how spatial and decisional coding evolved within a trial. In parallel to our fMRI analysis, we first computed the spatial encoding and recalibration indices for EEG activity patterns across time. Next, we used pattern component modelling to assess whether spatial and/or decisional codes arise at different latencies post-stimulus.

#### Spatial encoding and recalibration indices

To resolve spatial encoding across time we trained linear support vector regression models (SVR, LIBSVM^27^) on the EEG activity patterns of the pre-adaptation trials in overlapping 50 ms time windows sliding from −100 to 500 ms post-stimulus (EEG) and generalized them to trials from pre- and post-adaptation trials in a 4-fold cross-validation scheme. The cluster based bootstrap test on the Fisher z-transformed Pearson correlation coefficient between the true and the decoded sound locations revealed a significant cluster extending from 110 ms to 500 ms post-stimulus (p = 0.01 corrected for multiple comparisons within the entire [−50 - 500] ms window). The decoding accuracy was significantly better than chance from about 110 ms post-stimulus, rose steadily and peaked at about 355 ms (Figure 6A). Likewise, we assessed the recalibration index (RI) within the time window that showed a significant effect of spatial encoding, i.e., [110 - 500] ms post-stimulus. In the cluster-based bootstrap test the RI was significantly positive in two clusters from [185 - 285] ms (p = 0.019) and [335 - 470] ms (p = 0.005) post-stimulus (Figure 6B). Moreover, because previous ERP analyses suggested that the N100 potential is affected by recalibration^13^, we performed a temporal ROI analysis selectively on the EEG activity pattern averaged within the N100 time window (i.e., [70 - 130] ms). This temporal ROI analysis showed that the decoding accuracy was significantly better than chance (mean ± SEM: 0.125 ± 0.059; p = 0.0406) and the RI was significantly greater than zero (mean ± SEM: 5.963 ± 1.743; p = 0.0019).

**Figure 6.**
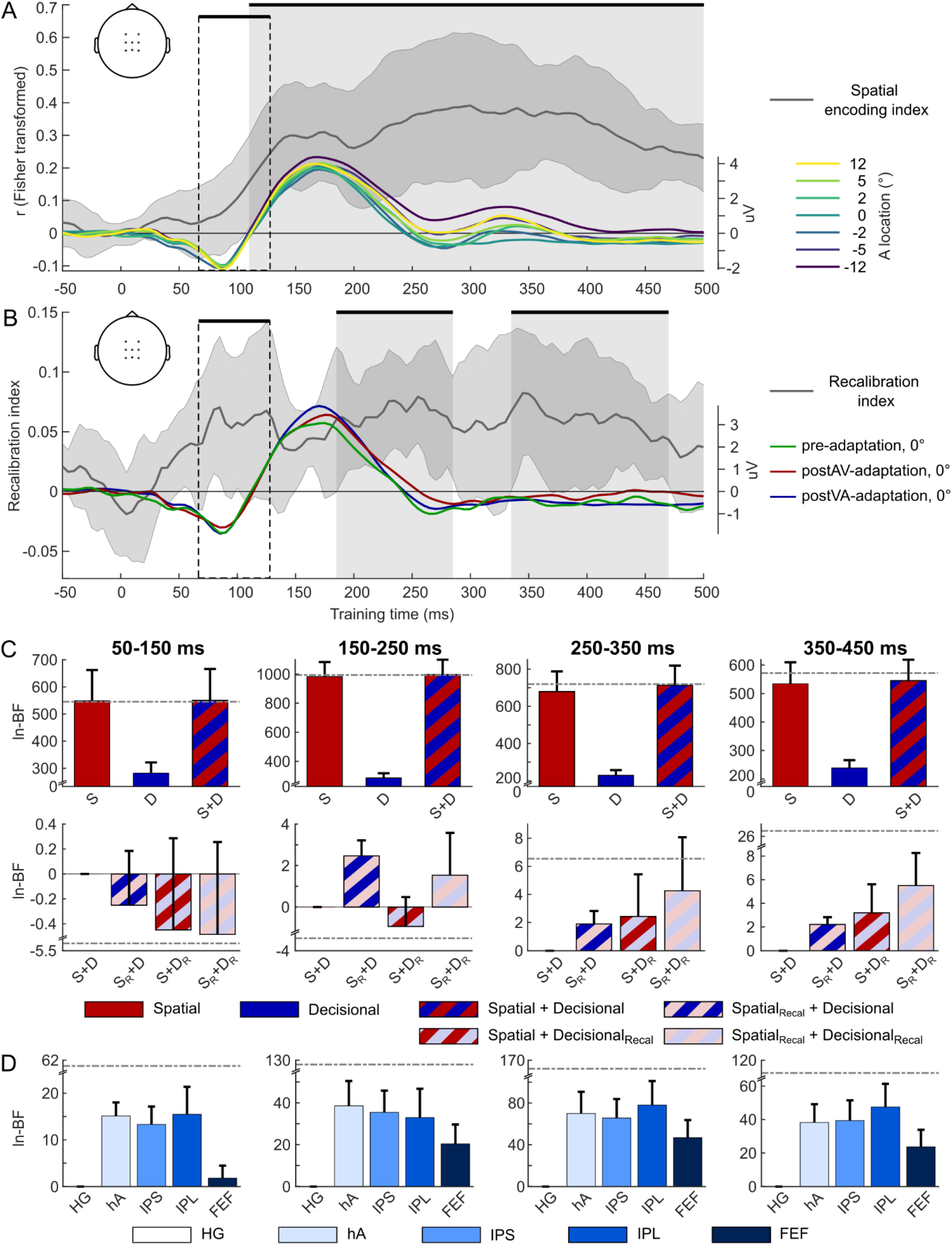
EEG results: multivariate decoding and pattern component modelling. **(A)** Time course of the spatial encoding index (i.e., across-subjects’ mean, dark grey line) and of EEG evoked potentials (across-subjects’ mean, averaged over central channels, see inset) for the 7 spatial locations in pre-adaptation phase (±12°, ±5°, ±2° and 0° azimuth). **(B)** Time course of the recalibration index (across-subjects’ mean, dark grey line) and the EEG evoked potentials (across-subjects’ mean, phases, averaged over central channels, see inset) for sound stimuli presented at 0° azimuth for pre-, postVA-, and postAV-adaptation. The shaded area around the spatial encoding index (A) and the recalibration index (B) indicates SEM. The shaded grey boxes and thick black horizontal lines (on top) indicate time windows in which the decoding accuracy (A) or the recalibration index (B) were significantly greater than zero at p < 0.05 corrected for multiple comparisons. Areas within the dashed boxes indicate the a priori defined time window [70 - 130] ms to focus on the early recalibration effects^13^. **(C)** PCM results – Spatial and/or decisional models as predictors for EEG activity patterns across four time windows. Across-subjects’ mean ln-BFs (± SEM) for each target model as indicated by the capital letter. (i) Top row: The spatial, decisional, and spatial + decisional models without recalibration relative to a null model that allows for no similarity between activity patterns. (ii) Bottom row: The models factorially manipulate whether the spatial and/or decisional component account for recalibration. Ln-Bayes factors are relative to the spatial + decisional model (without recalibration). Dash-dotted grey lines indicate the relative ln-BF for the fully flexible models as baseline. S = spatial model, D= decisional model, S_R_ = spatial model with recalibration, D_R_ = decisional model with recalibration. **(D)** PCM results – BOLD-response patterns from the five ROIs as predictors for EEG activity patterns across four time windows. Across-subjects’ mean ln-BFs (± SEM) for each target model relative to the model using BOLD-response patterns in HG as predictors. HG: Heschl’s gyrus; hA: higher auditory cortex; IPS: intraparietal sulcus; IPL: inferior parietal lobule; FEF: frontal eye field. ln-Bayes factors of > 1.1 are considered substantial evidence and > 3 as strong evidence for one model over the other^29^.

#### Model-based EEG analysis: dissociating perceptual and decisional codes

##### Pattern component modelling

Combining EEG and PCM we investigated whether spatial and decisional representations evolved with different time courses. For this, we compared the null, spatial, decisional, and combined spatial + decisional models as explanations for EEG activity patterns over spatial locations in four consecutive time windows: [50 - 150], [150 - 250], [250 - 350] and [350 - 450] ms post-stimulus (Figure 6C). From [50 - 150] ms the spatial model alone provided a sufficient explanation of the EEG data. From [150 - 250] ms and beyond the spatial + decisional model performed decisively better than the spatial model with ln-BFs of at least 10.9. These results suggest that the decisional code becomes progressively more important at later processing stages. This representational gradient across post-stimulus time was also apparent in our recalibration results (Figure 6C, bottom row). Spatial recalibration was expressed mainly via spatial coding (i.e., S_R_+D and S_R_+D_R_ better than S+D and S+D_R_ by ln-BFs of at least 2.4) from 150 to 250 ms and via decisional coding (i.e., S+D_R_ and S_R_+D_R_ better than S+D and S_R_+D by ln-BFs of at least 2.4) from 250 ms onwards. Nevertheless, as in our fMRI results recalibration relied jointly on spatial and decisional codes in later stages. Collectively, our EEG results revealed that spatial codes are more prominent in earlier processing stages and decisional coding in later ones. Critically, at later stages recalibration was expressed jointly in spatial and decisional codes.

##### Fusing EEG and fMRI in pattern component modelling

The fMRI results revealed a mixture of spatial and decisional codes in several ROIs across the dorsal processing hierarchy. To link fMRI and EEG results more directly, we fused them in PCMs in which the second moment matrices of the BOLD-response activation patterns in one of our five ROIs (i.e., HG, hA, IPS, IPL or FEF) formed the predictor for the distribution of EEG activity patterns over sound locations. Bayesian model comparison across these five fMRI-EEG fusion PCMs confirmed a direct relationship between our fMRI and EEG results (Figure 6D). The PCM with hA (i.e., including planum temporale) explained EEG activity patterns best (ln-BFs of at least 3.2) from [150 - 250] ms and the PCM with IPL from 250 ms onwards (ln-BFs of at least 8.0). We suspect that FEF has less explanatory power because of its smaller size, thereby contributing less to EEG scalp potentials. Collectively, combining EEG and fMRI spatiotemporally resolved the influence of recalibration on encoding of sound location and decisional uncertainty.

## Discussion

This study demonstrates that the brain recalibrates the senses by flexibly adapting spatial and decisional codes, which are expressed with opposite gradients along the auditory processing hierarchy. Early activity patterns in planum temporale encode sound locations in a continuous space that dynamically shifts towards misaligned visual inputs. Later activity patterns in frontoparietal cortices mainly code choice-related uncertainty in line with these representational changes.

Our behavioural results provide robust evidence that the brain recalibrates auditory space to keep auditory and visual maps in coregistration^15–17^. The point of subjective equality changed significantly between left and right recalibration in every observer (Figure 2). Because the audiovisual adaptation phase used a non-spatial task, this robust cross-sensory plasticity relied mainly on implicit perceptual rather than choice-related mechanisms^3,18,20,34–36^.

Consistent with previous research in non-human primates, our multivariate fMRI analyses showed that sound location can be decoded from activity patterns in a widespread network including primary auditory regions, planum temporale and frontoparietal cortices^7,19,24–26,37^. Moreover, the activity patterns in all of these regions adapted to changes in the sensory statistics as indicated by the neurometric functions and recalibration indices (Figure 3A and B). This widespread cross-sensory plasticity converges with our EEG results showing persistent spatial encoding and recalibration starting early with the auditory N1 component, generated in primary and secondary auditory cortices^38^, and extending until 500 ms post-stimulus (Figure 6A and B).

Critically, multidimensional scaling unravelled subtle representational differences in this network of regions (see Figure 3C). While recalibration induced a constant representational shift in planum temporale, a more complex similarity structure arose in IPL and FEF. We investigated whether this complex representational geometry resulted from multiplexing of spatial and decisional codes^39^ by comparing a spatial, a decisional, and a combined spatial + decisional model (Figure 4). The decisional model computes choice-related uncertainty based on the distance of the spatial estimates from the classification boundary (see Figure 4 and ^32^). The spatial hemifield model encodes sound location in the relative activity of two sub-populations of neurons each broadly tuned either to the ipsi- or contra-lateral hemifield^6–9^ - it is currently the leading model for spatial coding in human auditory cortices. Yet, we applied it as a model not only to auditory but also frontoparietal areas^7,25^, because its pattern similarity structure over the limited (i.e., [−12° to 12°]) sound locations is indistinguishable from that of the competing place code model (see Figure S1).

Crucially, Bayesian model comparison showed a double dissociation along the auditory processing stream: the spatial model provided a better explanation for the regional mean and fine-scale activation patterns in planum temporale, while the decisional model performed better in frontoparietal cortices. As shown in Figure 5A, the regional BOLD-response increased linearly for stimuli along the azimuth in planum temporale, but followed an inverted U-shaped function that adapts to audiovisual conflicts in frontoparietal cortices (Figure 5B top and bottom row)^7,19,24–26,40,41^. Likewise, the recalibration of the fine-scale activation patterns was better captured by the spatial model in planum temporale, but by the decisional model in frontoparietal cortices. Yet, despite this predominance of decisional coding, IPS, IPL and even FEF multiplexed spatial and decisional patterns (Figure 5C top row).

This spatial-decisional gradient across the cortical hierarchy was mirrored in their temporal evolutions. Recalibration of spatial coding was reflected in early EEG activity patterns ([150 - 250] ms), while adaptation of decisional uncertainty was more prominent in later activity (after 250 ms, Figure 6C bottom row). We could even directly link the double dissociations across time and regions by fusing fMRI and EEG via pattern component modelling: the BOLD-response patterns in hA predicted EEG activity patterns mainly at early ([150 - 250] ms) stages and IPL at later stages from 250 ms onwards (Figure 6D).

Collectively, our fMRI and EEG results demonstrate that the brain adapts flexibly to the changing statistics of the environment. They also highlight the critical importance to move beyond decoding analyses, as the ability to decode a feature value (e.g., sound location) from activity patterns does not imply that this feature is well represented^33^. Instead, in order to define the features or mixtures of features (e.g., spatial, decisional) that are encoded in activations, we need to characterize the fine-grained representational geometry and assess the explanatory power of different sets of features via model comparison. Only the latter was able to show that later recalibration effects in frontoparietal cortices reflect mainly choice-related rather than genuine spatial coding. The expression of recalibration in distinct spatial and decisional codes opens the intriguing possibility that recalibration may not always impact both codes together. Instead, the representations and neural circuitries involved in recalibration may depend on the particular task and/or the duration of recalibration. For instance, while our study focused on implicit recalibration using a non-spatial task, other studies have shown rapid recalibration effects when observers performed spatial localization tasks during the adaptation and/or post-adaptation phases^18,20,35,36^. It remains unknown whether these short-term recalibration effects involve plastic changes in the spatial representations or selectively alter choice-related activity. In line with this notion, accumulating research suggests that recalibration relies on multiple mechanisms that operate over different timescales^12,42^. Future studies thus need to disentangle the impact of recalibration duration and task on spatial and decisional coding.

So far, we have emphasized that spatial and decisional codes are expressed with opposite gradients across the cortical hierarchy and post-stimulus time. Yet, Bayesian model comparison also indicated that the combined spatial + decisional model significantly outperformed the more parsimonious decisional or spatial models throughout the cortical hierarchy. Likewise, a benefit was observed for the pattern component models in which recalibration was expressed in both spatial and decisional codes. These findings suggest that all higher order cortical regions (i.e., hA, IPS, IPL and FEF) multiplexed perceptual and decisional codes and adapt both to recalibrate the senses^39^. Moreover, because multiplexing of spatial and decisional codes arose mainly later from 250 ms post-stimulus, the coding of decisional uncertainty in auditory cortices (hA) is likely to reflect top-down influences from frontoparietal areas^43,44^.

In conclusion, our results show that audiovisual adaptation relies on spatial and decisional coding. Crucially, the expression of spatial and decisional codes evolves with different time courses and opposite gradients along the auditory processing hierarchy. Early neural activity in planum temporale encoded sound location within a continuous auditory space. Later frontoparietal activity mainly encoded observers’ decisional uncertainty that flexibly adapts in accordance with these transformations of auditory space.

## Methods

### Participants

15 participants (10 females, mean age = 22.1; SD = 4.1) participated in the psychophysics study. Five of those participants (4 females, mean age = 22.2; SD = 5.1, one author of the study, A.M.) completed the fMRI and EEG experiments (for full selection criteria see Supplementary methods). All participants had no history of neurological or psychiatric illnesses, normal or corrected-to-normal vision and normal hearing. They gave informed written consent to participate in the study, which was approved by the research ethics committee of the University of Birmingham (approval number: ERN_11_0470AP4) and conducted in accordance with the Declaration of Helsinki.

### Stimuli

The auditory stimulus consisted of a burst of white noise with a duration of 50 ms and 5 ms on/off ramp delivered at a 75 dB sound pressure level. To create a virtual auditory spatial signal, the noise was convolved with spatially specific head-related transfer functions thereby providing both binaural and monaural cues for sound location^45^. Head-related transfer functions from the available locations in the MIT database (http://sound.media.mit.edu/resources/KEMAR.html) were interpolated to the desired spatial locations. For the EEG experiment, scanner background noise was superimposed on the spatial sound stimuli to match the task environment of the fMRI experiment. The visual stimulus was a cloud of 15 white dots (diameter = 0.4° visual angle) sampled from a bivariate Gaussian distribution with a vertical and horizontal standard deviation of 1.5° and a duration of 50 ms presented on a dark grey background (90% contrast) in synchrony with the auditory stimuli.

### Experimental design and procedure

The psychophysics, fMRI, and EEG experiments used the same design including three phases: (i) unisensory auditory pre-adaptation, (ii) audiovisual adaptation (AV- or VA-adaptation) and (iii) unisensory auditory post-adaptation (postAV-adaptation or postVA-adaptation) (Figure 1B). Psychophysics, fMRI, EEG included two days each for left (i.e., VA) and two days for right (i.e., AV) adaptation (except for one participant who completed EEG over two days), i.e., 12 days of experimental testing + 1 day pre-screening for each participant. The order of left and right adaptation was counterbalanced across participants. Throughout all experiments, participants fixated a central fixation cross (0.5° diameter).

#### Auditory pre- and post-adaptation

In unisensory pre- and post-adaptation phases (Figure 1B), participants were presented with auditory stimuli that were sampled uniformly from 7 spatial locations (±12°, ±5°, ±2° and 0° visual angle) along the azimuth (stimulus onset asynchrony (SOA) = 2000 ms ± 200 ms jitter). Participants performed a two-alternative forced-choice left-right spatial classification task explicitly only on a fraction of trials (22%), the so-called ‘response trials’, which were randomly interspersed and indicated 500 ms after sound onset by a brief (i.e., 200 ms duration) dimming of the fixation cross to 55% of its initial contrast. The fMRI decoding was based only on the non-response trials to minimize motor confounds.

Participants indicated their left-right spatial classification response by pressing one of two buttons with their index or middle fingers of their left or right hand. The response hand was alternated over runs within a day to control for potential motor confounds (see fMRI multivariate decoding section below). The order of left and right response hands was counter-balanced across days (for number of trials, runs etc., see Supplementary methods).

#### Audiovisual adaptation

In the audiovisual adaptation phase, participants were presented with spatially disparate (±15° visual angle) audiovisual stimuli (SOA = 500 ms): the visual stimulus was uniformly sampled from three locations along the azimuth (i.e., −5°, 0°, 5°). On separate days, the visual stimulus was spatially shifted by 15° either to the left (i.e., VA-adaptation) or to the right (i.e., AV-adaptation) with respect to the auditory stimulus. Hence, we included the following audiovisual stimulus location pairs: [A = −20°, V = −5°], [A = −15°, V = 0°], [A = −10°, V = 5°] in (right) AV-adaptation phases and [A = 10°, V = −5°], [A = 15°, V = 0°], [A = 20°, V = 5°] in (left) VA-adaptation phases. The locations of the audiovisual stimulus pairs were fixed within mini-blocks of 5 (psychophysics, i.e., duration of 2.5 s) or 20 (fMRI, EEG, i.e., duration of 10 s) consecutive trials.

Participants detected slightly dimmer visual stimuli (80% of normal contrast), pseudorandomly interspersed on 10% of the trials (i.e., so-called ‘response trials’). This non-spatial task ensures maintenance of participants’ attention and introduces audiovisual recalibration at the perceptual rather than decisional or motor response level. To allow sufficient time for responding (given the short SOA of 500 ms), we ensured that each response trial was followed by at least 3 consecutive non-response trials (see Supplementary methods for further details).

### Experimental setup

In all experiments, visual and auditory stimuli were presented using Psychtoolbox version 3.0.11^46,47^ under MATLAB R2011b (MathWorks Inc.) on a MacBook Pro running Mac OSX 10.6.8 (Apple Inc.). In the psychophysics and EEG experiments, participants were seated at a desk with their head rested on a chinrest. Two accessory rods were mounted on the chin rest serving as forehead rest and allowing stable and reliable head positioning. Visual stimuli were presented at a viewing distance of 60 cm via a gamma-corrected 24’’ LCD monitor (ProLite B2483HS, iiyama Corp.) with a resolution of 1920 × 1080 pixels at a frame rate of 60 Hz. Auditory stimuli were delivered via circumaural headphones (HD 280 Pro, Sennheiser electronic GmbH & Co. KG) in the psychophysics experiment and via in-ear earphones (E-A-RTONE GOLD, 3M Company Auditory Systems) in the EEG experiment. Participants used a standard USB keyboard for responding. In the fMRI experiment, visual stimuli were back projected to a plexiglass screen using a D-ILA projector (DLA-SX21, JVC, JVCKENWOOD UK Ltd.) with a resolution of 1400 × 1050 pixels at a frame rate of 60 Hz. The screen was visible to the subject through a mirror mounted on the magnetic resonance (MR) head coil and the eye-to-screen distance was 68 cm. Auditory stimuli were delivered via a pair of MR compatible headphones (MR Confon HP-VS03, Cambridge Research Systems Ltd). Participants responded using a two-button MR-compatible keypad (LXPAD 1×5-10M, NATA Technologies). Exact audiovisual onset timing in adaptation trials was confirmed by recording visual and auditory signals concurrently with a photodiode and a microphone.

### Eye movement recording

Eye movement recordings were calibrated in the recommended field of view (32° horizontally and 24° vertically) for the EyeLink 1000 Plus system (SR Research Ltd.) with the desktop mount at a sampling rate of 2000 Hz. Eye position data were on-line parsed into events (saccade, fixation, eye blink) using the EyeLink 1000 Plus software. The ‘cognitive configuration’ was used for saccade detection (velocity threshold = 30°/sec, acceleration threshold = 8000°/sec^2^, motion threshold = 0.15°) with an additional criterion of radial amplitude > 1°. Fixation position was post-hoc offset corrected. In the fMRI experiment, precise positioning of participants’ heads inside the scanner bore was critical for the sensitive measurement of spatial recalibration, so that high quality eye movement recordings were not possible.

### Behavioural analysis for psychophysics, fMRI, and EEG experiments

#### Signal detection measures

Participants responded only on a fraction of ‘response trials’, i.e., 22% in auditory pre- and post-adaptation and 10% in AV- and VA-adaptation. This enabled us to assess their performance with the signal sensitivity measure d’:

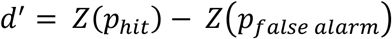

where *p_hit_* and *P_false alarm_* are the hit and false alarm rates, respectively. Hits are ‘response trials’, on which observers gave a response. False alarms are ‘non-response trials’, on which observers gave a response. 100% hit rate and 0% false alarm rate were approximated by 99.999% and 0.001%, respectively, to enable the calculation of Z-scores.

#### Psychometric functions

For the pre-, postVA- and postAV-adaptation phases we fitted cumulative Gaussians as psychometric functions (PF) to the percentage ‘perceived right’ responses on the ‘response trials’ as a function of stimulus location (±12°, ±5°, ±2° and 0°) (see Figure 2, and Supplementary methods). The models were fitted individually to the behavioural data of each participant with maximum-likelihood estimation as implemented in the Palamedes toolbox^48^ (www.palamedestoolbox.org). To enable reliable parameter estimates for each participant, we employed a multi-condition fitting using the following constraints: (i) the just noticeable differences (JND or slope parameter) were set equal across all conditions (i.e., pre-, postVA- and postAV-adaptation phases); (ii) guess and lapse rates were set equal to each other and (iii) equal across all conditions. Furthermore, (iv) we constrained the fitted guess and lapse rate parameters to be within 0 and 0.1.

For statistical inference, we assessed the goodness of fit of the cumulative Gaussian for each condition and participant using a likelihood ratio test. This likelihood ratio test compares the likelihood of participants’ responses given the model that is constrained by a cumulative Gaussian function (i.e., our ‘target model’) to the likelihood given by a so-called ‘saturated model’ that models observers’ responses with one parameter for each stimulus location in each condition. The resulting likelihood ratio for the original data set is then compared with a null distribution of likelihood ratios generated by parametrically bootstrapping the data (5000x) from the ‘target model’ fitted to the original data set and refitting the ‘target’ and ‘saturated’ models. Since the likelihood ratio for the original data set was not smaller in any of the participants than 5% of the parametrically bootstrapped likelihood ratios (i.e., p > 0.05), we inferred sufficient goodness of fit for all participants.

Next, we assessed whether AV- and VA-adaptation induced a shift in participants’ perceived auditory location by comparing a ‘static’ model, which constrains PSEs to be equal for pre-, postVA- and postAV-adaptation, with a ‘recalibration’ model, which includes three PSE values for the pre-, postVA-, and postAV-adaptation PFs. For each participant and model, we calculated the Akaike Information Criterion (AIC^28^) as an approximation to the model evidence. We performed Bayesian model comparison at the random effects level as implemented in SPM12 to obtain the protected exceedance probability for the candidate models^49^.

### fMRI data acquisition and analysis

#### fMRI data acquisition

We used a 3T Philips Achieva scanner to acquire both T1-weighted anatomical images (TR/TE/TI, 7.4/3.5/min. 989 ms; 176 slices; image matrix, 256 × 256; spatial resolution, 1 × 1 × 1 mm^3^ voxels) and T2*-weighted echo-planar images (EPI) with blood oxygenation level-dependent (BOLD) contrast (fast field echo; TR/TE, 2800/40 ms; 38 axial slices acquired in ascending direction; image matrix, 76 × 75; slice thickness, 2.5 mm; interslice gap, 0.5 mm; spatial resolution, 3 × 3 × 3 mm^3^ voxels). On each of the four days we acquired five auditory pre-adaptation runs and 4 audiovisual adaptation runs. Each fMRI run started and ended with 10 s fixation and the first 4 volumes of each run were discarded to allow for T1 equilibration effects.

#### Pre-processing and general linear model

The data were analysed with Statistical Parametric Mapping (SPM12; http://www.fil.ion.ucl.ac.uk/spm/^50^). Scans from each participant were realigned using the first as a reference, unwarped and slice-time corrected. The time series in each voxel was high-pass filtered to 1/128 Hz. The EPI images were spatially smoothed with a Gaussian kernel of 3 mm FWHM and analysed in native space. The data were modelled in a mixed event/block fashion with regressors entered into the design matrix after convolving the unit impulse or the block with a canonical hemodynamic response function and its first temporal derivative. In the unisensory auditory pre- and post-adaptation phases, unisensory sound stimuli were modelled as events separately for each of our 7 (sound location) × 2 (response vs. non-response) × 3 (pre, post-VA, post-AV) conditions. In the AV- and VA-adaptation phases, AV stimulus presentations were modelled as blocks separately for the 3 (visual locations) × 2 (VA vs. AV adaptation) conditions. In addition, we modelled all response trials during the adaptation phase with a single regressor to account for motor responses. Realignment parameters were included as nuisance covariates. Condition-specific effects for each subject were estimated according to the general linear model (GLM). To minimise confounds of motor response, we limited all subsequent fMRI analyses to the parameter estimates pertaining to the ‘non-response’ trials.

For the BOLD-response analysis, we computed contrast images comparing auditory stimulus at a particular location > fixation in each subject (averaged over runs) resulting in 21 contrast images (i.e., 7 (sound location) × 3 (pre, postVA, postAV)). Moreover, we computed a contrast and associated t-image that compared all 21 sound conditions relative to fixation baseline (for identification of sound-responsive voxels).

For the multivariate decoding and representational similarity analyses, we applied multivariate spatial noise normalization to the parameter estimates using the noise covariance matrix obtained from the residuals of the GLM and the optimal shrinkage method^51^ and finally performed Euclidean normalization.

#### Regions of interest for fMRI analysis

We defined five regions of interest (ROI, combined from two hemispheres) that have previously been implicated in auditory spatial processing based on neurophysiology and neuroimaging research^10,24,25^. Heschl’s gyrus (HG), higher auditory cortex (hA) and inferior parietal lobule (IPL) were defined using the following parcellations of the Destrieux atlas of Freesurfer 5.3.0^52^: (i) HG: Heschl’s gyrus and anterior transverse temporal gyrus; (ii) hA: higher auditory cortex, i.e., transverse temporal sulcus, planum temporale and posterior ramus of the lateral sulcus; (iii) IPL: inferior parietal lobule, i.e., supramarginal gyrus and inferior part of the postcentral sulcus. The intraparietal sulcus (IPS) and frontal eye field (FEF) were defined using the following group-level retinotopic probabilistic maps^53^: (iv) IPS: IPS0, IPS1, IPS2, IPS3, IPS4, IPS5 and SPL1; (v) FEF: hFEF. All probabilistic maps were thresholded to a probability of 0.1 (i.e., probability that a vertex belongs to a particular ROI) and inverse normalized into each participant’s native space.

#### fMRI multivariate decoding – spatial encoding and recalibration indices

We extracted the voxel response patterns in a particular ROI from the pre-whitened and normalized parameter estimate images pertaining to the magnitude of the BOLD-response for each condition and run. To avoid motor confounds, we used the parameter estimate images only from the ‘non-response trials’. In a 4-fold stratified cross-validation procedure, we trained support vector regression models (C = 1, υ = 0.5, LIBSVM 3.17^27^) to learn the mapping from the condition-specific fMRI response patterns (i.e., examples) to external spatial locations (i.e., labels) using examples selectively from the unisensory auditory pre-adaptation runs of all but one fold^54,55^. This learnt mapping was used to decode the spatial locations from the BOLD-response patterns of the remaining pre-adaptation fold and all postVA- and postAV-adaptation examples (acquired in separate runs).

To determine whether a ROI encodes auditory spatial representations, we computed the Pearson correlation coefficients between the true and the decoded auditory locations for the pre-adaptation runs for each participant as a ‘spatial encoding index’.

To determine whether auditory spatial representations in a region of interest are recalibrated by misaligned visual signals, we binarized the predicted auditory locations into left vs. right predictions and computed the difference in the fraction of ‘decoded right responses’ between auditory postVA- and postAV-adaptation phases as ‘recalibration index’ (RI).

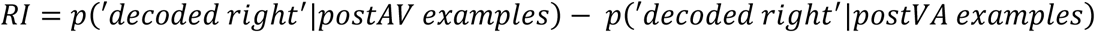

Importantly, postVA- and postAV-adaptation phases were matched in terms of time, exposure, training, and other non-specific effects.

To allow for generalization at the population level, we entered the subject-specific Fisher z-transformed spatial encoding and recalibration indices into separate bootstrap-based one sample t-tests against zero at the group level^56^. Briefly, an empirical null distribution of t-values was generated from the data by resampling the subject-specific values with replacement, subtracting the across-subjects mean and computing the one-sample t-statistic for each bootstrap sample. A right-tailed p-value was computed as the proportion of t-values in the bootstrapped t-distribution greater than the observed t-value.

#### fMRI multivariate decoding – neurometric functions

We fitted cumulative Gaussians as neurometric functions (NF^57^) in each ROI to the percentage of ‘decoded right’ averaged across participants as a function of stimulus location. Consistent with our behavioural analysis we assessed whether AV- and VA-adaptation induced a shift in the decoded location of the unisensory auditory stimuli by comparing a ‘static’ model with a ‘recalibration’ model using the Akaike Information Criterion (AIC^28^, for details see Supplementary methods).

#### fMRI multivariate pattern analysis – representational dissimilarity analyses and multidimensional scaling

We generated 21 condition specific contrast images for the 7 auditory spatial locations × 3 (pre-, postVA-, and postAV-adaptation) by averaging parameter estimate images across fMRI runs for each participant. We then characterized the geometry of spatial representations using representational dissimilarity matrices (RDMs^30^) based on the Mahalanobis distance for each participant and each ROI separately for pre-adaptation as well as postVA- and postAV-adaptation phases (see Supplementary methods). Using non-classical multidimensional scaling (MDS) with non-metric scaling, we projected the group level RDMs (i.e., averaged across participants) onto a one-dimensional space (‘reflecting’ spatial dimension along the azimuth).

### EEG data acquisition and analysis

#### EEG data acquisition

Continuous EEG signals were recorded from 64 channels using Ag/AgCl active electrodes arranged in 10-20 layout (ActiCap, Brain Products GmbH, Gilching, Germany) at a sampling rate of 1000 Hz with FCz as reference. Channel impedances were kept below 10 kΩ.

#### EEG pre-processing

Pre-processing was performed with the FieldTrip toolbox^58^ (http://www.fieldtriptoolbox.org/). Raw data were high pass filtered at 0.1 Hz, inspected for bad channels, re-referenced to the average of all channels, and low pass filtered at 45 Hz. Bad channels were identified by visual inspection, rejected and interpolated using the neighbouring channels. Trial epochs for the unisensory auditory pre-adaptation and the auditory post-adaptation conditions were extracted between [−100 - 500] ms post-stimulus (i.e., the onset of the response cue on the response trials), baseline corrected and down-sampled to 200 Hz. Epochs containing artefacts within the time window of interest (i.e., between [0 - 500] ms post-stimulus) were identified based on visual inspection and rejected. Furthermore, based on eyetracking data, trials were rejected if they (i) contained eye blinks or (ii) saccades or (iii) the eye gaze was away from the fixation cross by more than 2 degrees (% rejected trials across-subjects mean ± SEM: 8.2 ± 1.0%). Grand average ERPs were computed by averaging all trials for each condition first within each participant and then across participants.

For the multivariate analysis, we applied spatial multivariate noise normalization to the individual trials using a noise covariance matrix estimated separately for each time point and the optimal shrinkage method^51^. Furthermore, the EEG activity patterns were divided by their Euclidean norm separately for each time point and trial for normalization. Since the EEG responses on trials with and without behavioural response were identical until 500 ms post-stimulus, we pooled over response and no response trials in all EEG analyses.

#### EEG multivariate decoding – spatial encoding and recalibration indices

Similar to our fMRI analysis, we trained a SVR in a 4-fold stratified cross-validation (C = 1, υ = 0.5^27^) to learn the mapping from evoked potentials (averages of 16 randomly sampled trials) of the pre-adaptation run to external auditory space over 50 ms time windows, shifting in increments of 5 ms, from −100 ms to 500 ms post-stimulus. The learnt mapping was used to decode the sound location from the EEG activity patterns of the pre-adaptation examples in the remaining fold and all post-adaptation examples. To minimize sampling variance, we averaged the decoded locations across 50 repetitions of this cross-validation procedure to compute the ‘spatial encoding’ and ‘recalibration’ indices for each 50 ms window. At the random effects level, we report results from a bootstrap-based t-test^56^ against zero corrected for multiple comparisons across [−50 - 500] ms using a cluster-based correction with an auxiliary cluster-defining height threshold of p < 0.05 uncorrected^59^.

Based on our a priori hypothesis that spatial location is encoded in the N100 potential^13^, we also performed bootstrap-based t-tests on the spatial encoding and recalibration indices obtained from evoked potentials averaged within a [70 - 130] ms window.

### Spatial hemifield and decisional models for fMRI and EEG

We assessed a spatial and a decisional encoding model as explanations for the regional mean BOLD-response and fine-scale fMRI/EEG patterns.

#### Spatial hemifield model

The hemifield model^6–9^ encodes sound location in the relative activity of two sub-populations of neurons each broadly tuned either to the ipsi- or contra-lateral hemifield (Figure 4A). We simulated 360 neurons with broad Gaussian tuning functions. The standard deviation was set to 64°. The means of the tuning functions were sampled uniformly from 80° to 100° for the neuronal population tuned to the contralateral hemifield and from −80° to −100° azimuth for the neuronal population tuned to the ipsilateral hemifield. Consistent with previous research^8^ the ratio of the ipsi- and contralaterally tuned neurons was set to 30%/70% (see Figure 4A, S1A).

For the pre-adaptation conditions we sampled neural responses from the seven sound locations in our paradigm. For the post-adaptation conditions, we sampled again from these seven locations for the non-recalibration model. For the recalibration model we sampled the neural responses from the above locations shifted by 2.3° to the right (postVA-adaptation) or left (postAV-adaptation). The shift by 2.3° was calculated as the difference between the across-subjects mean PSE values in postVA- and postAV-adaptation phases from the psychometric functions (for comparison of spatial hemifield and place code models see Supplementary methods).

#### Decisional model

In the decisional model the activity of a neuron encodes observers’ choice-related uncertainty that depends non-linearly on the distance between observers’ spatial estimates and their left-right spatial classification boundary^32^ according to

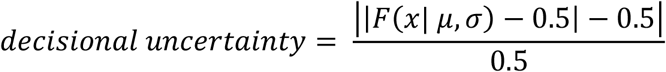

where *F*(*x*| *μ,σ*) is the cumulative normal distribution function with mean *μ* and standard deviation *σ* evaluated at spatial location *x:*

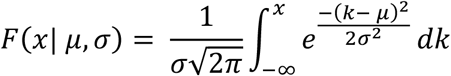

The standard deviations of the cumulative normal distribution were set to 10° and their means were uniformly sampled between −1° and +1° (see Figure 4A). We simulated responses from 360 neurons. Exactly as for the spatial model, we sampled neural responses from the seven sound locations for the pre-adaptation conditions and the post-adaptation conditions for the non-recalibration model. For the recalibration model we sampled the neural responses from the above locations shifted by 2.3° to the right (postVA-adaptation) or left (postAV-adaptation).

As shown in Figure 4B and C, the spatial and decisional models make distinct predictions for the regional mean BOLD-response and the pattern similarity structure over the 21 conditions = 7 spatial locations × 3 phases (pre, postVA, postAV).

##### Regional mean BOLD-response: spatial and decisional linear mixed effects models

###### Regional mean BOLD-response

For each of the 2 (hemisphere: left, right) × 5 (ROI: HG, hA, IPS, IPL, FEF) regions we selected the 20 most reliably responsive voxels, i.e., with the greatest t-values for all unisensory sound conditions relative to fixation. For each of those 10 regions we extracted the BOLD-response magnitude for each of the 7 locations × 3 phases (pre-, postAV-, and postVA-adaptation) and formed the regional mean.

###### Linear mixed effects modelling

To account for lateralization effects^7^, we performed separate analyses for each region and hemisphere. Separately for each hemisphere we used the spatial (resp. decisional) model to generate predicted regional mean BOLD-responses for each of the 21 conditions = 7 locations × 3 phases (pre-, postAV-, and postVA-adaptation) by averaging activations of 360 simulated neurons.

We generated seven linear mixed effects (LME) models that varied in their fixed effects predictors:

- Null LME: single intercept term.
- Spatial LME model (S): predictor from the spatial encoding model without recalibration and intercept term.
- Decisional LME model (D): predictor from the decisional model without recalibration and intercept term.

The remaining LME models included spatial, decisional and intercept terms (i.e., 3 fixed effects regressors) and factorially manipulated whether the spatial and/or the decisional predictor modelled recalibration:

- (S+D) Spatial without recalibration + decisional without recalibration
- (S_R_+D) Spatial with recalibration + decisional without recalibration,
- (S+D_R_) Spatial with recalibration + decisional without recalibration
- (S_R_+D_R_) Spatial with recalibration + decisional with recalibration

Subject level effects were included as random effects. For each of the 2 hemispheres × 5 ROIs we fitted these seven LME models using maximum likelihood estimation and computed the Bayesian information criterion (BIC). The figures show the natural logarithm of the Bayes factors (ln-Bayes factors) averaged across hemispheres for LME models relative to the null LME (for visualization details, see Supplementary methods).

##### Multivariate pattern: pattern component modelling of fMRI and EEG data

To assess whether spatial and/or decisional models can explain the fine-scale fMRI or EEG activity patterns across the 7 (sound location) × 3 (pre, post-VA, post-AV) = 21 conditions, we combined Pattern Component Modelling (PCM^33^, https://github.com/jdiedrichsen/pcm_toolbox) and Bayesian model comparison. Like the more widely used representational similarity analyses, pattern component modelling allows one to investigate whether specific representational structures – as implied for instance by the spatial or decisional model – are expressed in fMRI or EEG activity patterns. Critically, PCM goes beyond standard representational similarity analyses by allowing us to assess arbitrary mixtures of representational components such as a combined spatial + decisional model (i.e., with unknown mixture weights, for further information see Supplementary methods).

Consistent with our LME analysis, we generated second moment matrices (‘pattern components’) as predictors for PCM based on the activations of 360 simulated neurons from the spatial and decisional models, respectively.

We compared the following PCM models for fMRI and EEG data:

*Null PCM:* all conditions are independent (i.e., the 2^nd^ moment matrix is the identity matrix).

*Spatial PCM* (S): activity patterns generated by the spatial model without recalibration. *Decisional PCM* (D): activity patterns generated by the decisional model without recalibration.

*Combined (spatial + decisional) PCMs:* activity patterns are a weighted linear combination of the patterns generated by the spatial and the decisional model. We factorially manipulated whether the spatial and/or decisional model accommodate audiovisual recalibration:

- (S+D) spatial component without recalibration and decisional component without recalibration
- (S_R_+D) spatial component with recalibration and decisional component without recalibration
- (S+D_R_) spatial component without recalibration and decisional component with recalibration
- (S_R_+D_R_) spatial component with recalibration and decisional component with recalibration

*Free (i.e., fully flexible) PCM* imposes no constraints on the second moment matrix and provides an upper benchmark. If a model performs at least as good as the cross-validated free PCM, it is sufficiently complex to capture all consistent variation in the data^33^.

###### fMRI and EEG fusion PCMs

To assess whether the fMRI activation patterns evolve with different time courses in EEG, we computed five PCMs each including only one pattern component generated from the BOLD-response patterns of HG, hA, IPS, IPL and FEF.

###### Model estimation and comparison

In our fMRI analysis these PCM models were applied to the pre-whitened parameter estimates from the first level GLM analysis separately for each of the 5 fMRI ROIs (i.e., HG, hA, IPS, IPL, FEF pooled over both hemispheres). In our EEG analysis they were applied to pre-whitened evoked EEG potentials averaged within individual runs (20 trials) and within each of the 4 EEG time windows (i.e., [50 - 150] ms, [150 - 250] ms, [250 - 350] ms, [350 - 450] ms). We estimated the parameters of the PCM models in a leave-one-subject-out cross-validation scheme at the group level^33^. Because the parameter estimates were computed relative to common baseline, we modelled the run mean as a fixed effect in fMRI when comparing the spatial, decisional, and spatial + decisional models without recalibration. When comparing the four spatial + decisional models with/without recalibration, we modelled the run effects as random, because a fixed effect run mean would have modelled out the recalibration effect across runs. In EEG, we modelled the run effect as a random effect (as they were not computed relative to a common baseline).

The marginal likelihood for each model and subject from the leave-one-subject-out cross-validation scheme was used as an approximation to the model evidence. We compared the models using the natural logarithm (ln) of the Bayes factors averaged across participants^33^, with ln-Bayes factors of > 1.1 = substantial evidence and > 3 = strong evidence^29^.

## Acknowledgements

This research was funded by the European Research Council (https://erc.europa.eu/; grant number: ERC-2012-StG_20111109 multsens).

## Author Contributions

Conceptualization: M.A., A.M., and U.N.; Methodology: M.A. and A.M.; Software: M.A. and A.M.; Formal Analysis: M.A., A.M., and U.N.; Investigation: M.A. and A.M.; Resources: U.N.; Data Curation: M.A. and A.M.; Writing - Original Draft: M.A., A.M., and U.N.; Writing - Review & Editing: M.A., A.M., and U.N.; Visualization: M.A. and A.M.; Supervision: U.N.; Funding Acquisition: U.N.

## Declaration of Interests

The authors have declared that no competing interests exist.

## Notes

### Competing Interest Statement

The authors have declared no competing interest.

### Summary of Updates

Streamlined Methods section; Updated Supplemental information

## References

1. Chen, L. & Vroomen, J. Intersensory binding across space and time: a tutorial review. Atten. Percept. Psychophys. 75, 790–811 (2013).

2. Grothe, B., Pecka, M. & McAlpine, D. Mechanisms of sound localization in mammals. Physiol. Rev. 90, 983–1012 (2010).

3. Kopčo, N., Lin, I.-F., Shinn-Cunningham, B. G. & Groh, J. M. Reference frame of the ventriloquism aftereffect. J. Neurosci. 29, 13809–13814 (2009).

4. Maier, J. X. & Groh, J. M. Multisensory guidance of orienting behavior. Hear. Res. 258, 106–12 (2009).

5. Wandell, B. A., Dumoulin, S. O. & Brewer, A. A. Visual field maps in human cortex. Neuron 56, 366–383 (2007).

6. McAlpine, D., Jiang, D. & Palmer, A. R. A neural code for low-frequency sound localization in mammals. Nat. Neurosci. 4, 396–401 (2001).

7. Ortiz-Rios, M. et al. Widespread and Opponent fMRI Signals Represent Sound Location in Macaque Auditory Cortex. Neuron 93, 971–983 (2017).

8. Salminen, N. H., May, P. J. C., Alku, P. & Tiitinen, H. A population rate code of auditory space in the human cortex. PLOS ONE 4, e7600 (2009).

9. Stecker, G. C., Harrington, I. A. & Middlebrooks, J. C. Location coding by opponent neural populations in the auditory cortex. PLoS Biol. 3, 0520–0528 (2005).

10. Schlack, A., Sterbing-D’Angelo S. J., Hartung, K., Hoffmann, K.-P. & Bremmer, F. Multisensory Space Representations in the Macaque Ventral Intraparietal Area. J. Neurosci. 25, 4616–4625 (2005).

11. Bertelson, P., Frissen, I., Vroomen, J. & de Gelder, B. The aftereffects of ventriloquism: patterns of spatial generalization. Percept. Psychophys. 68, 428–436 (2006).

12. Bosen, A. K., Fleming, J. T., Allen, P. D., O’Neill, W. E. & Paige, G. D. Multiple time scales of the ventriloquism aftereffect. PLOS ONE 13, e0200930 (2018).

13. Bruns, P., Liebnau, R. & Röder, B. Cross-modal training induces changes in spatial representations early in the auditory processing pathway. Psychol. Sci. 22, 1120–6 (2011).

14. Frissen, I., Vroomen, J., de Gelder, B. & Bertelson, P. The aftereffects of ventriloquism: generalization across sound-frequencies. Acta Psychol. (Amst.) 118, 93–100 (2005).

15. Radeau, M. & Bertelson, P. The after-effects of ventriloquism. Q. J. Exp. Psychol. 26, 63–71 (1974).

16. Recanzone, G. H. Rapidly induced auditory plasticity: the ventriloquism aftereffect. Proc. Natl. Acad. Sci. U. S. A. 95, 869–875 (1998).

17. Woods, T. M. & Recanzone, G. H. Visually induced plasticity of auditory spatial perception in macaques. Curr. Biol. 14, 1559–64 (2004).

18. Wozny, D. R. & Shams, L. Recalibration of auditory space following milliseconds of cross-modal discrepancy. J. Neurosci. 31, 4607–12 (2011).

19. Zierul, B., Röder, B., Tempelmann, C., Bruns, P. & Noesselt, T. The role of auditory cortex in the spatial ventriloquism aftereffect. NeuroImage 162, 257–268 (2017).

20. Park, H. & Kayser, C. Shared neural underpinnings of multisensory integration and trial-by-trial perceptual recalibration in humans. eLife 8, (2019).

21. Zwiers, M. P., Van Opstal, A. J. & Paige, G. D. Plasticity in human sound localization induced by compressed spatial vision. Nat. Neurosci. 6, 175–181 (2003).

22. Mullette-Gillman, O. A., Cohen, Y. E. & Groh, J. M. Eye-centered, head-centered, and complex coding of visual and auditory targets in the intraparietal sulcus. J. Neurophysiol. 94, 2331–2352 (2005).

23. Michalka, S. W., Rosen, M. L., Kong, L., Shinn-Cunningham, B. G. & Somers, D. C. Auditory Spatial Coding Flexibly Recruits Anterior, but Not Posterior, Visuotopic Parietal Cortex. Cereb. Cortex 26, 1302–1308 (2016).

24. Rauschecker, J. P. & Tian, B. Mechanisms and streams for processing of ‘what’ and ‘where’ in auditory cortex. Proc. Natl. Acad. Sci. U. S. A. 97, 11800–11806 (2000).

25. van der Heijden, K., Rauschecker, J. P., de Gelder, B. & Formisano, E. Cortical mechanisms of spatial hearing. Nat. Rev. Neurosci. 20, 609–623 (2019).

26. Zatorre, R. J., Bouffard, M., Ahad, P. & Belin, P. Where is ‘where’ in the human auditory cortex? Nat. Neurosci. 5, 905–9 (2002).

27. Chang, C.-C. & Lin, C.-J. LIBSVM: a library for support vector machines. ACM Trans. Intell. Syst. Technol. 2, 1–27 (2011).

28. Akaike, H. A New Look at the Statistical Model Identification. in Selected Papers of Hirotugu Akaike (eds. Parzen, E., Tanabe, K. & Kitagawa, G.) 215–222 (Springer New York, 1998). doi:10.1007/978-1-4612-1694-0_16.

29. Jeffreys, H. Theory of probability. (Oxford University Press, 1961).

30. Nili, H. et al. A toolbox for representational similarity analysis. PLOS Comput. Biol. 10, e1003553 (2014).

31. Middlebrooks, J. C., Clock, A. E., Xu, L. & Green, D. M. A panoramic code for sound location by cortical neurons. Science 264, 842–844 (1994).

32. Grinband, J., Hirsch, J. & Ferrera, V. P. A Neural Representation of Categorization Uncertainty in the Human Brain. Neuron 49, 757–763 (2006).

33. Diedrichsen, J., Yokoi, A. & Arbuckle, S. A. Pattern component modeling: A flexible approach for understanding the representational structure of brain activity patterns. NeuroImage 180, 119–133 (2018).

34. Zaidel, A., Ma, W. J. & Angelaki, D. E. Supervised calibration relies on the multisensory percept. Neuron 80, 1544–1557 (2013).

35. Bruns, P. & Röder, B. Sensory recalibration integrates information from the immediate and the cumulative past. Sci. Rep. 5, 12739 (2015).

36. Mendonça, C., Escher, A., van de Par, S. & Colonius, H. Predicting auditory space calibration from recent multisensory experience. Exp. Brain Res. 233, 1983–1991 (2015).

37. Tian, B., Reser, D., Durham, A., Kustov, A. & Rauschecker, J. P. Functional specialization in rhesus monkey auditory cortex. Science 292, 290–293 (2001).

38. Winkler, I., Denham, S. & Escera, C. Auditory Event-related Potentials. in *Encyclopedia of Computational Neuroscience* (eds. Jaeger, D. & Jung, R.) 1–29 (Springer, 2013). doi:10.1007/978-1-4614-7320-6_99-1.

39. Raposo, D., Kaufman, M. T. & Churchland, A. K. A category-free neural population supports evolving demands during decision-making. Nat. Neurosci. 17, 1784–1792 (2014).

40. Petkov, C. I., Kayser, C., Augath, M. & Logothetis, N. K. Functional Imaging Reveals Numerous Fields in the Monkey Auditory Cortex. PLOS Biol. 4, e215 (2006).

41. Zimmer, U. & Macaluso, E. High Binaural Coherence Determines Successful Sound Localization and Increased Activity in Posterior Auditory Areas. Neuron 47, 893–905 (2005).

42. Watson, D. M., Akeroyd, M. A., Roach, N. W. & Webb, B. S. Distinct mechanisms govern recalibration to audio-visual discrepancies in remote and recent history. Sci. Rep. 9, 8513 (2019).

43. Mihalik, A. & Noppeney, U. Causal inference in audiovisual perception. J. Neurosci. (2020) doi:10.1523/JNEUROSCI.0051-20.2020.

44. Werner, S. & Noppeney, U. Distinct functional contributions of primary sensory and association areas to audiovisual integration in object categorization. J. Neurosci. 30, 2662–2675 (2010).

45. Gardner, W. G. & Martin, K. D. HRTF measurements of a KEMAR. J. Acoust. Soc. Am. 97, 3907–3908 (1995).

46. Brainard, D. H. The Psychophysics Toolbox. Spat. Vis. 10, 433–436 (1997).

47. Kleiner, M., Brainard, D. & Pelli, D. What’s new in Psychtoolbox-3? in Perception, 36 (EVCP Abstract Supplement) (2007).

48. Prins, N. & Kingdom, F. A. A. Applying the Model-Comparison Approach to Test Specific Research Hypotheses in Psychophysical Research Using the Palamedes Toolbox. Front. Psychol. 9, (2018).

49. Rigoux, L., Stephan, K. E., Friston, K. J. & Daunizeau, J. Bayesian model selection for group studies - Revisited. NeuroImage 84, 971–985 (2014).

50. Friston, K. J. et al. Statistical parametric maps in functional imaging: A general linear approach. Hum. Brain Mapp. 2, 189–210 (1994).

51. Ledoit, O. & Wolf, M. A well-conditioned estimator for large-dimensional covariance matrices. J. Multivar. Anal. 88, 365–411 (2004).

52. Destrieux, C., Fischl, B., Dale, A. & Halgren, E. Automatic parcellation of human cortical gyri and sulci using standard anatomical nomenclature. NeuroImage 53, 1–15 (2010).

53. Wang, L., Mruczek, R. E. B., Arcaro, M. J. & Kastner, S. Probabilistic Maps of Visual Topography in Human Cortex. Cereb. Cortex N. Y. N 1991 25, 3911–3931 (2015).

54. Rohe, T. & Noppeney, U. Distinct computational principles govern multisensory integration in primary sensory and association cortices. Curr. Biol. 1, 1–6 (2016).

55. Rohe, T. & Noppeney, U. Cortical hierarchies perform Bayesian causal inference in multisensory perception. PLOS Biol. 13, e1002073 (2015).

56. Efron, B. & Tibshirani, R. J. An Introduction to the bootstrap. (Springer US, 1993). doi:10.1007/978-1-4899-4541-9.

57. Rohe, T. & Noppeney, U. Reliability-Weighted Integration of Audiovisual Signals Can Be Modulated by Top-down Attention. eNeuro 5, ENEURO.0315-17.2018 (2018).

58. Oostenveld, R., Fries, P., Maris, E. & Schoffelen, J.-M. FieldTrip: Open source software for advanced analysis of MEG, EEG, and invasive electrophysiological data. Comput. Intell. Neurosci. 2011, 156869 (2011).

59. Maris, E. & Oostenveld, R. Nonparametric statistical testing of EEG- and MEG-data. J. Neurosci. Methods 164, 177–190 (2007).

