## Supplemental Information for "Audiovisual adaptation is expressed in spatial and decisional codes"

### Supplementary Information

#### Supplementary Results

##### Eye movement results

To control for eye-movement-related confounds, we recorded participants' eye movements during the psychophysics and EEG experiments. In the fMRI experiment, precise positioning of participants' head inside the scanner bore was critical for the sensitive measurement of spatial recalibration, which did not allow high quality eye-movement recordings. As EEG recordings are highly susceptible to eye-movement-related confounds in spatial tasks [1], we rejected all trials in the EEG experiment that did not fulfil stringent criteria (% rejected trials across subjects mean ± SEM: 8.2 ± 1.0%; see EEG Pre-processing section for criteria).

In the psychophysics experiment fixation was well maintained throughout the entire experiment with participants fixating correctly on 94.02% ± 1.00% (mean ± SEM) of all trials (i.e., eye gaze was maintained within a radius of 2° and without saccades or eyeblinks) and post-stimulus saccades detected on only 1.24% ± 0.33% (mean ± SEM) of all trials (including response and non-response trials in all pre-, postAV/postVA- and AV-adaptation phases). In one-way repeated measures ANOVAs we assessed whether eye movement indices (i.e., % saccades, % eye blinks, and post-stimulus mean horizontal fixation position) differed across the 3 experimental phases (pre-, postAV-, and postVA-adaptation) in the non-response trials (i.e., the trials that were included in our fMRI analysis). No significant differences were found across the 3 experimental phases for % saccades and % blinks, however, we observed a significant effect of experimental phase on the mean horizontal eye position ($F(2,14) = 8.02$, $p = 0.002$). Fisher's LSD post-hoc tests revealed significant differences only between pre- and post-adaptation phases (mean difference between pre- and postVA-adaptation = 0.061°, $p = 0.004$; mean difference between pre- and postAV-adaptation = 0.047°, $p = 0.027$). Importantly, we did not observe a significant difference in the mean horizontal fixation position between postAV- and postVA-adaptation phase ($p = 0.147$). In other words, the direction of recalibration did not affect observers' eye position. This is critical because we assess the effects of recalibration by directly comparing trials from postAV- and postVA-adaptation phases to avoid temporal confounds.

#### Supplementary methods

##### Participants

Nineteen right-handed participants took part in the pre-screening session. 15 of those participants (10 females, mean age = 22.1; SD = 4.1) returned for the psychophysics part of the study. Participants were included after the pre-screening session if (i) they provided reliable spatial discrimination responses in the auditory pre-adaptation phase, i.e., the just noticeable difference (JND) of the psychometric function fitted to the spatial discrimination responses was < 4°; (ii) reliably discriminated between response and non-response trials, i.e., sensitivity index (d') > 2.5 and (iii) reliably detected the target luminance change during the audiovisual adaptation phase, i.e., d' > 2.5 (see Experiment design and procedure for details). Six of the psychophysics participants that had the highest recalibration effect size were selected to participate in the fMRI and EEG experiments (see [2]). Five of those participants (4 females, mean age = 22.2; SD = 5.1) completed the whole study (one was an author of the study, A.M.). One participant was excluded after 3 fMRI days because their sensitivity index d' during the adaptation phase was ≤ 2.5 (see above iii). All participants had no history of neurological or psychiatric illnesses, normal or corrected-to-normal vision and normal hearing. All participants gave informed written consent to participate in the study, which was approved by the research ethics committee of the University of Birmingham (approval number: ERN_11_0470AP4) and was conducted in accordance with the principles outlined in the Declaration of Helsinki.

#### Auditory pre- and post-adaptation

To increase design efficiency (in the fMRI study), stimuli were presented from one of the 7 possible locations in a pseudorandomized order in ~36 s blocks of 18 stimuli interleaved with 6 s fixation. In the pre-adaptation phase one run consisted of 10 blocks (i.e., 18 stimuli x 10 blocks = 180 stimuli) and 9 fixation intervals and lasted for ~7 minutes. After five blocks we inserted a longer fixation interval of 15 s. In the post-adaptation phase, each run started with an audiovisual adaptation block (see next section) followed by 5 post-adaptation blocks. In total, a run included 2 audiovisual adaptation and 10 post-adaptation blocks (~14 minutes). As in the pre-adaptation phase, the post-adaptation blocks were interleaved with 6 s fixation.

The EEG experiment followed the same overall structure as the fMRI experiment, except that the fixation intervals were not included to reduce the duration of the experiment. In the psychophysics experiment, runs in the pre-adaptation phase consisted of 5 stimulus blocks (i.e., 18 stimuli x 5 blocks = 90 stimuli) and 4 fixation intervals (~3.5 minutes).

To maintain participants' engagement with the task, they received visual feedback after every 5 pre- or post-adaptation blocks (i.e., 90 trials) about their detection performance (i.e., the dimming of the fixation cross, n.b. not their localization performance) based on the sensitivity index *d'* (i.e., d' > 3.5: smiling face, 2.5 < d' < 3.5: neutral face, 2.5 < d': sad face).

The pre-adaptation phase of the fMRI (resp. EEG) experiment included 3600 auditory trials (2800 no response + 800 response trials, i.e., 18 stimuli per block x 10 blocks per run x 5 runs per day x 4 days). The postVA (resp. postAV)-adaptation phase included 1440 auditory trials (1120 no response + 320 response trials) for postAV-adaptation and the same number of trials for postVA-adaptation phase (i.e., 18 stimuli per block x 10 blocks per run x 4 runs per day x 2 days). In the psychophysics experiment, the pre-adaptation phase included 1800 auditory trials (1400 no response + 400 response trials, i.e., 18 stimuli per block x 5 blocks per run x 5 runs per day x 4 days). The postVA (resp. postAV)-adaptation phase included 1800 auditory trials (1400 no response + 400 response trials) for postAV-adaptation and the same number of trials for postVA-adaptation phase (i.e., 18 trials per block x 10 blocks per run x 5 runs per day x 2 days)

#### Audiovisual adaptation

One audiovisual adaptation block included either 360 trials (for the first adaptation block in psychophysics and typically in fMRI and EEG, i.e., 3 minutes) or 120 trials (for the remaining adaptation blocks in psychophysics, i.e., 1 minute). To maintain participants' engagement with the task, they received visual feedback after every adaptation block (i.e., 360 or 120 trials) about their detection performance (i.e. of the dimmer visual stimuli) based on the sensitivity index *d'* (i.e., d' > 3.5: smiling face, 2.5 < d' < 3.5: neutral face, 2.5 < d': sad face).

In total, both fMRI and EEG studies included 5760 audiovisual adaptation trials (5184 no response + 576 response trials) for the (left) VA-adaptation phase (i.e., 360 trials per block x 2 blocks per run x 4 runs per day x 2 days). Likewise, there were 5760 audiovisual trials for the (right) VA-adaptation phase.

##### Organization of each testing day

Each day started with 5 pre-adaptation runs amounting to 900 trials per day (i.e. 18 trials per block x 10 stimulation blocks per run x 5 runs). The subsequent 4 runs (resp. 5 runs in psychophysics) included 2 VA (or AV)-adaptation blocks and 10 postVA (or postAV)-adaptation blocks, such that each VA (or AV)-adaptation block was followed by 5 postVA (or postAV)-adaptation blocks. Hence, each day included 2880 trials AV adaptation trials (i.e, 360 per block x 2 blocks per run x 4 runs per day in fMRI and EEG and 2400 AV adaptation trials in psychophysics). Further, each day included 720 post-adaptation trials in fMRI and EEG (i.e. 18 trials per block x 10 blocks per run x 4 runs per day) and 900 trials in psychophysics (because of 5 runs per day, see Figure 1B). The AV adaptation and post-adaptation blocks were interleaved with 15 s of fixation.

#### fMRI data acquisition and analysis

##### fMRI multivariate decoding – neurometric functions

Consistent with our behavioural analysis we assessed spatial encoding and recalibration in our ROIs using neurometric functions (NF[3]). For this, we first binarized the decoded sound locations into ‘decoded left’ and ‘decoded right’ and computed the percentage of ‘decoded right’ for each sound location and participant. To enable reliable parameter estimates, we employed multi-condition fitting using the following constraints: (i) the JNDs (i.e., slope parameter) were constrained to be equal across all conditions; (ii) guess and lapse rates were set to be equal to each other and (iii) constrained to be equal across all conditions; (iv) lapse rates were set to be within 0 and 0.45 (to account for the lower signal-to-noise ratio of fMRI data). Likelihood ratio test as described in the behavioural analysis section confirmed adequate goodness of fit for the neurometric functions in each ROI (i.e., $p>0.05$ across all ROIs, Table S3). We assessed whether AV- and VA-adaptation induced a shift in the decoded location of the unisensory auditory stimuli. Consistent with our behavioural analysis, we compared a ‘static or no recalibration’ model with a ‘recalibration’ model. The static model assumes no effect of audiovisual spatial adaptation on unisensory auditory spatial representations by constraining the point of subjective equality (PSE) for the pre-, postVA-, and postAV-adaptation PFs to be equal. The adaptation model includes three PSE values for the pre-, postVA-, and post-AV-adaptation PFs thereby accommodating potential shifts in the PSE values as a result of audiovisual recalibration.

##### fMRI multivariate pattern analysis – representational dissimilarity analyses and multidimensional scaling

We characterized the spatial representations using representational similarity analysis (RSA, as implemented in the rsatoolbox[4]. For pre-adaptation, we computed a rank-transformed (and scaled between 0 and 1) 7 x 7 RDM based on the 7 auditory contrast images from the pre-adaptation phase. To characterize and visualize the effect of AV- and VA-adaptation on the representational geometry of spatial representations we computed a 14 x 14 matrix (i.e., 7 spatial locations for postVA-adaptation and 7 spatial locations for postAV-adaptation) where we rank-transformed (and scaled between 0 and 1) the 7 x 7 sub-components of the post-adaptation RDMs (i.e., separately for the postAV- and postVA-adaptation). For visualization (see Figure 3C) we averaged the rank transformed matrices across participants.

#### Comparison of spatial hemifield and place code models

While accumulating research has shown that sound location is coded by broadly tuned neural populations in auditory cortices in mammals[5,6], less is known about auditory spatial coding in parietal or frontal cortices from single cell neurophysiology. For instance, one previous neurophysiology study in non-human primates has suggested that neurons in parietal area VIP have narrow auditory receptive fields that are congruent with visual receptive fields[7]. Critically, the current study selected only near-central sound locations to optimize the characterization of recalibration rather than to dissociate between different spatial encoding principles. As we will show in the simulations below, the hemifield model and a place code model with narrow tuning functions make indistinguishable predictions for the pattern similarity structure over those near-central locations. We have therefore used the hemifield model as a generic perceptual spatial encoding model and note that comparable predictions could be made with the place code model.

In simulations, we investigated how the proportion of neurons tuned to different locations influences the pattern similarity structure. For the hemifield model we simulated 360 neurons with broad Gaussian tuning functions. The standard deviation was set to 64˚ as in our main study. The means of the tuning functions were sampled uniformly from 80˚ to 100 ˚ for the neuronal population tuned to the contralateral hemifield and from -80 ˚ to -100 ˚ azimuth for the neuronal population tuned to the ipsilateral hemifield. To illustrate how the ratio of ipsi- vs contralaterally tuned neurons affect the predictions of the model we performed two simulations with different ipsi/contra ratios: 30%/70% as suggested by previous research[8] and 50%/50%.

For the spatial model based on the place code, we simulated 360 neurons with narrow Gaussian tuning functions (SD = 26˚) with means distributed between -179˚ and 180˚ azimuth. The means were distributed either uniformly in 1˚ increments or non-uniformly, with most of the neurons responding to the contralateral hemifield (see Figure S1A).

As in our main analysis, for the pre-adaptation conditions we sampled neural responses from the seven sound locations (i.e., -12˚, -5˚, -2˚, 0˚, 2˚, 5˚, 12˚) in our paradigm. For the post-adaptation conditions, we sampled again from these seven locations for the non-recalibration model. For the recalibration model we sampled the neural responses from the above locations shifted by 2.3˚ to the right (postVA-adapation) or left (postAV-adaptation). The shift by 2.3˚ was calculated as the difference between the across-subjects mean PSE values in postVA- and postAV-adaptation phases from the psychometric functions.

As shown in Figure S1A, the hemifield models with ipsi/contra ratios of 30%/70% and 50%/50% make different predictions for the regional mean BOLD-response, but not for the pattern similarity structure over locations. Likewise, the place code models with uniform and non-uniform sampling of neurons with different tuning functions differ in their predictions for the regional mean BOLD-response, but not for the pattern similarity structure. In both cases, i.e., unequal ratios and non-uniform sampling can introduce an increase in the regional BOLD-response for sounds presented along the azimuth. By contrast, the similarity structure is relatively immune to these changes in model parameters.

#### Plotting of regional mean BOLD-responses (Figure 5A)

For visualization purposes only (i.e., Figure 5A), we averaged the BOLD-responses across hemispheres whilst controlling for the fact that the effects of spatial location and recalibration differ between the hemispheres as indicated by our model comparison results[9]. For the pre-adaptation conditions, we averaged the BOLD-responses for [-12°, -5°, -2°, 0°, 2°, 5°, 12°] in the left hemisphere with [12°, 5°, 2°, 0°, -2°, -5°, -12°] in the right hemisphere. For the post-adaptation conditions, we averaged the BOLD-responses for [-12°, -5°, -2°, 0°, 2°, 5°, 12°] of the postVA-adaptation conditions in the left hemisphere with [12°, 5°, 2°, 0°, -2°, -5°, -12°] of the postAV-adaptation conditions in the right hemisphere. In this way, we obtained estimates of the BOLD-response magnitude for each of the 21 conditions for a representative left hemisphere region across all five ROIs (Figure 5A).

#### Multivariate pattern: pattern component modelling of fMRI and EEG data

PCM estimates the likelihood of the observed neural activity under a multivariate linear model:

$$\boldsymbol{Y} = \boldsymbol{ZU} + \boldsymbol{XB} + \mathbf{E}$$

$$\boldsymbol{u_p} \sim \boldsymbol{N}(\mathbf{0}, \boldsymbol{G})$$

$$\boldsymbol{\varepsilon_p} \sim \boldsymbol{N}(\mathbf{0}, \boldsymbol{I\sigma}^{\mathbf{2}})$$

$\boldsymbol{Y}$ is an (N x P) data matrix. N is equal to the number of conditions (i.e., 21) times the number of runs; P is equal to the number of measurement channels (fMRI voxels or EEG channels). Each row in $\boldsymbol{Y}$ includes the pre-whitened activity pattern, i.e., across-voxel parameter estimate patterns from the first-level GLM in fMRI and across-electrode evoked potentials in EEG. $\boldsymbol{Z}$ is a (N x 21 conditions) design matrix specifying the experimental conditions with 21 columns (i.e., 21 conditions = 7 sound locations x [pre, postVA, postAV]). $\boldsymbol{U}$ is a matrix of true activity profiles (21 conditions x number of channels). $\boldsymbol{X}$ is the design matrix for the patterns of no interest $\boldsymbol{B}$ (e.g., condition-independent means of each voxel-specific activity profile). $\mathbf{E}$ is the N x P matrix with the error terms. PCM treats the true activity profiles $\boldsymbol{u_p}$ of each measurement channel (the columns of $\boldsymbol{U}$) as random variables sampled from a multivariate normal distribution $\boldsymbol{u_p} \sim \boldsymbol{N}(\mathbf{0}, \boldsymbol{G})$**.** Here, $\boldsymbol{G}$ is the second moment matrix of activity profiles, which describes the similarity structure over the 21 conditions in the experiment.

$\boldsymbol{G}$ can be expressed as a linear combination of second moment matrices (e.g., generated from the decisional and/or spatial model) with $\exp(\theta_h)$ determining the weight of each second moment component to the second moment matrix $\boldsymbol{G}$ estimated from the measured neural activity profiles over conditions.

$$G = \sum_h \exp(\theta_h) G_h$$

PCM estimates the marginal likelihood of the observed data given the model and the model parameters $\boldsymbol{\theta}$ (i.e., the weights for the different second moment component matrices, signal strength, noise variance) by integrating over the actual activity profiles:

$$p(\boldsymbol{Y}|\boldsymbol{\theta}) = \int p(\boldsymbol{Y}|\boldsymbol{U}, \boldsymbol{\theta}) p(\boldsymbol{U}|\boldsymbol{\theta}) d\boldsymbol{U}$$

Hence, the estimation of the individual activity profiles is not necessary and the second moment matrix $\boldsymbol{G}$ (i.e., in our study a 21 x 21 matrix) is a sufficient statistic to compute the marginal likelihoods [10].

#### Supplementary Tables

**Table S1. Behavioural results: hit rate, d' and PSE in the sound localization and adaptation tasks.**

| | Experiment | Hit rate | d' | PSE |
|---|---|---|---|---|
| **Pre-adaptation phase** | | | | |
| | Psychophysics | 93.1% (±1.1%) | 4.08 (±0.19) | 0.59 (±0.22) |
| | fMRI | 97.5% (±1.0%) | 4.80 (±0.29) | -0.72 (±0.48) |
| | EEG | 91.3% (±2.7%) | 4.09 (±0.20) | -0.34 (±0.21) |
| **Adaptation phase** | | | | |
| AV-adaptation | Psychophysics | 96.4% (±1.4%) | 4.65 (±0.21) | not applicable |
| | fMRI | 92.4% (±1.1%) | 4.54 (±0.08) | |
| | EEG | 89.6% (±4.0%) | 4.38 (±0.53) | |
| VA-adaptation | Psychophysics | 95.6% (±1.6%) | 4.69 (±0.22) | |
| | fMRI | 89.5% (±2.7%) | 4.09 (±0.23) | |
| | EEG | 88.1% (±2.8%) | 4.21 (±0.26) | |
| **Post-adaptation phase** | | | | |
| postAV-adaptation | Psychophysics | 91.4% (±1.7%) | 4.07 (±0.20) | -0.95 (±0.29) |
| | fMRI | 97.8% (±0.6%) | 5.01 (±0.23) | -3.34 (±0.21) |
| | EEG | 90.8% (±3.4%) | 4.53 (±0.21) | -2.33 (±0.33) |
| postVA-adaptation | Psychophysics | 91.2% (±1.4%) | 4.09 (±0.19) | 2.10 (±0.25) |
| | fMRI | 98.1% (±0.6%) | 5.18 (±0.30) | 1.66 (±0.54) |
| | EEG | 88.8% (±4.3%) | 4.31 (±0.37) | 1.87 (±0.41) |

All behavioural results are reported as across subjects' mean (± SEM). Number of subjects was 15 in psychophysics and 5 in fMRI and EEG experiments. The percentage of responses made when a response trial was indicated (Hit rate) and the corresponding sensitivity index (d') was computed separately for each experiment (Psychophysics, fMRI, and EEG) and experimental phases (pre-adaptation, adaptation and postAV/postVA-adaptation). In pre-and post-adaptation phases only, psychometric functions were fitted to the sound localization results. The point of subjective equality (PSE) values are reported based on the 'recalibration' model (see Methods). SEM: standard error of the mean.

**Table S2. Psychometric functions for psychophysics, fMRI, and EEG experiments: Bayesian model comparison of the 'static model' and the 'recalibration model'**

| **Model** | | **Static model** | **Recalibration model** |
|---|---|---|---|
| | Number of parameters | 3 | 5 |
| **Psychophysics** | AIC | -4641.7 | -5212.4 |
| | Exp. post. prob. | 0.0588 | 0.9412 |
| | Exceedance prob. | 0 | 1 |
| | Prot. exceedance prob. | 0.0003 | 0.9997 |
| **fMRI** | AIC | -2143.1 | -2509.1 |
| | Exp. post. prob. | 0.1104 | 0.8896 |
| | Exceedance prob. | 0.0002 | 0.9998 |
| | Prot. exceedance prob. | 0.1124 | 0.8876 |
| **EEG** | AIC | -1721.7 | -2030.1 |
| | Exp. post. prob. | 0.1104 | 0.8896 |
| | Exceedance prob. | 0.0002 | 0.9998 |
| | Prot. exceedance prob. | 0.1124 | 0.8876 |

The static model assumes that the psychometric function is identical for the pre-, postVA-, and postAV-adaption phases (i.e., three parameter estimates: 1 PSE, 1 JND and 1 guess/lapse rate). The recalibration model accommodates the possibility that the point of subjective equality (PSE) shifts as a result of recalibration by including separate PSEs for the pre-, postAV-, and postVA-adaptation phases as free parameters (in total 5 parameters: 3 PSEs, 1 JND and 1 guess/lapse rate). AIC: Akaike Information Criterion at the group level (subject level AICs summed up over subjects, AIC = LL - n; LL = natural logarithm (ln) of the likelihood, n = number of parameters). More positive AIC indicates a better model. AIC difference > 1.1 substantial evidence[11]. Exp. post. prob. = expected posterior probability; Exceedance prob. = exceedance probability; Prot. exceedance prob. = protected exceedance probability.

**Table S3. Neurometric functions for fMRI: Bayesian model comparison of the 'static model' and the 'recalibration model'**

| Statistic \ ROI | HG | hA | IPS | IPL | FEF |
|---|---|---|---|---|---|
| Goodness of fit test | p = 0.8858 | p = 0.3836 | p = 0.2852 | p = 0.1274 | p = 0.8548 |
| AIC (Static model) | -860.6 | -690.9 | -721.8 | -807.6 | -837.1 |
| AIC (Recalibration model) | -847.7 | -670.7 | -704.8 | -797.3 | -831.4 |

Goodness of fit is assessed by comparing the loglikelihood ratio of the target relative to a saturated model for the original data relative to a bootstrapped null distribution of loglikelihood ratios. Sufficient goodness of fit is inferred if $p > 0.05$ (see Methods). The static model assumes that the psychometric function is identical for the pre-, postVA-, and postAV-adaptation phases (i.e., 3 parameter estimates: 1 PSE, 1 JND and 1 guess/lapse rate). The recalibration model accommodates the possibility that the point of subjective equality (PSE) shifts as a result of recalibration by including separate PSEs for the pre-, postAV-, and postVA-adapation phases as free parameters (in total 5 parameters: 3 PSEs, 1 JND and 1 guess/lapse rate). AIC: Akaike Information Criterion (AIC = LL - n; LL = natural logarithm (ln) of the likelihood, n = number of parameters). More positive AIC indicates a better model. AIC difference > 1.1 considered substantial evidence [11].

**Table S4. fMRI regional BOLD-response: Spatial and decisional Linear mixed effects models**

| ROI / Model | HG | hA | IPS | IPL | FEF |
|---|---|---|---|---|---|
| S | -1.5 | 4.4 | -1.1 | -1.3 | -0.3 |
| D | -2.3 | -1.1 | 3.2 | -0.2 | 0.3 |
| S+D | -3.8 | 2.9 | 2.4 | -1.5 | 0.4 |
| $S_R$+D | -3.7 | 3.8 | 1.8 | -2.2 | -0.2 |
| S+$D_R$ | -3.7 | 2.4 | 7.2 | 1.7 | 5.5 |
| $S_R$+$D_R$ | -3.7 | 3.2 | 6.6 | 1.1 | 4.9 |

ln-Bayes factors (relative to the null model) for the linear mixed effects (LME) models fitted to the regional BOLD-response separately for HG, hA, IPS, IPL, FEF relative to a null model. The target models included predictors from the decisional, spatial, and decisional + spatial models with and without recalibration. The null model includes only a constant. ln-Bayes factors > 1.1 are considered substantial evidence[11]. S: spatial model without recalibration, D: decisional model without recalibration, S+D: spatial model without recalibration + decisional model without recalibration, $S_R$+D: spatial model with recalibration + decisional model without recalibration, S+$D_R$: spatial model without recalibration + decisional model with recalibration, $S_R$+$D_R$: spatial model with recalibration + decisional model with recalibration null: null model, ln: natural logarithm, LME: linear mixed effects, ROI: region of interest, HG: Heschl's gyrus, hA: higher auditory cortex, IPS: intraparietal sulcus, IPL: inferior parietal lobule, FEF: frontal eye field.

**Table S5. fMRI activation pattern: Pattern component modelling**

| Model \ ROI | HG | hA | IPS | IPL | FEF |
|---|---|---|---|---|---|
| S (rel null) | 8.0 +/- 4.7 | 119.0 +/- 29.1 | 135.2 +/- 86.6 | 29.5 +/- 10.2 | 7.6 +/- 6.2 |
| D (rel null) | -0.7 +/- 0.7 | -5.9 +/- 3.6 | -21.3 +/- 10.3 | 10.1 +/- 7.5 | 9.0 +/- 4.4 |
| S+D (rel null) | 8.3 +/- 4.8 | 142.1 +/- 36.9 | 190.0 +/- 110.1 | 65.0 +/- 27.7 | 37.7 +/- 20.9 |
| $S_R$+D (rel S+D) | 0.5 +/- 0.3 | 3.9 +/- 1.7 | 3.4 +/- 2.1 | 1.1 +/- 0.7 | 0.8 +/- 0.6 |
| S+$D_R$ (rel S+D) | -0.2 +/- 0.3 | 1.2 +/- 0.6 | 22.2 +/- 9.2 | 9.6 +/- 3.3 | 15.1 +/- 3.8 |
| $S_R$+$D_R$ (rel S+D) | 0.4 +/- 0.4 | 5.1 +/- 1.6 | 25.9 +/- 10.8 | 11.0 +/- 4.1 | 16.1 +/- 4.3 |

Across subjects' mean ± SEM of the relative ln-Bayes factors for each pattern component model (relative to its particular baseline model as indicated in parentheses) fitted to the fMRI activation patterns separately for HG, hA, IPS, IPL, FEF. (i) the spatial, decisional, and spatial + decisional PCMs relative to a null model that assumes activity patterns over all sound locations are independent. (ii) the spatial + decisional PCMs that factorially manipulate whether recalibration is incorporated in the spatial and/or decisional components relative to the spatial + decisional model without recalibration. ln-Bayes factors > 1.1 considered substantial evidence[11]. S: spatial model without recalibration, D: decisional model without recalibration, S+D: spatial model without recalibration + decisional model without recalibration, $S_R$+D: spatial model with recalibration + decisional model without recalibration, S+$D_R$: spatial model without recalibration + decisional model with recalibration, $S_R$+$D_R$: spatial model with recalibration + decisional model with recalibration, Free: free model, null: null model, rel: relative to, ln: natural logarithm, PCM: pattern component modelling, ROI: region of interest, HG: Heschl's gyrus, hA: higher auditory cortex, IPS: intraparietal sulcus, IPL: inferior parietal lobule, FEF: frontal eye field.

**Table S6. EEG PCM analysis ln-Bayes factors**

| Model \ Time win | 50-150 ms | 150-250 ms | 250-350 ms | 350-450 ms |
|---|---|---|---|---|
| S (rel null) | 549.2 +/- 112.9 | 982.0 +/- 99.8 | 678.8 +/- 108.3 | 534.2 +/- 75.1 |
| D (rel null) | 282.4 +/- 39.5 | 287.3 +/- 32.0 | 229.3 +/- 27.9 | 239.2 +/- 27.1 |
| S+D (rel null) | 550.5 +/- 115.1 | 996.6 +/- 101.5 | 712.4 +/- 105.6 | 545.1 +/- 73.3 |
| $S_R$+D (rel S+D) | -0.3 +/- 0.4 | 2.5 +/- 0.7 | 1.9 +/- 0.9 | 2.2 +/- 0.6 |
| S+$D_R$ (rel S+D) | -0.4 +/- 0.7 | -0.9 +/- 1.4 | 2.4 +/- 3.0 | 3.2 +/- 2.4 |
| $S_R$+$D_R$ (rel S+D) | -0.5 +/- 0.7 | 1.5 +/- 2.0 | 4.3 +/- 3.8 | 5.5 +/- 2.8 |
| hA (rel HG) | 15.1 +/- 2.9 | 38.5 +/- 11.9 | 70.0 +/- 20.7 | 38.2 +/- 10.9 |
| IPS (rel HG) | 13.2 +/- 3.8 | 35.3 +/- 10.4 | 65.6 +/- 18.2 | 39.4 +/- 12.1 |
| IPL (rel HG) | 15.4 +/- 5.9 | 32.8 +/- 13.7 | 78.0 +/- 23.0 | 47.5 +/- 13.8 |
| FEF (rel HG) | 1.8 +/- 2.7 | 20.4 +/- 9.3 | 46.8 +/- 16.8 | 23.7 +/- 10.2 |

Across subjects' mean ± SEM of the relative ln-Bayes factors for each pattern component model (relative to its particular baseline model as indicated in parentheses) fitted to the EEG activity patterns separately for four time windows. (i) the spatial, decisional, and spatial + decisional PCMs relative to a null model that assumes activity patterns over all sound locations are independent. (ii) the spatial + decisional PCMs that factorially manipulate whether recalibration is incorporated in the spatial and/or decisional components relative to the spatial + decisional model without recalibration. (iii) the PCMs including pattern components generated from the BOLD-response patterns of one of the four ROIs (combined over both hemispheres) relative to the HG model: hA, IPS, IPL and FEF. ln-Bayes factors > 1.1 considered substantial evidence [11]. S: spatial model without recalibration, D: decisional model without recalibration, S+D: spatial model without recalibration + decisional model without recalibration, $S_R$+D: spatial model with recalibration + decisional model without recalibration, S+$D_R$: spatial model without recalibration + decisional model with recalibration, $S_R$+$D_R$: spatial model with recalibration + decisional model with recalibration, Free: free model, null: null model, rel: relative to, ln: natural logarithm, PCM: pattern component modelling, ROI: region of interest, HG: Heschl's gyrus, hA: higher auditory cortex, IPS: intraparietal sulcus, IPL: inferior parietal lobule, FEF: frontal eye field.

#### Supplementary figures

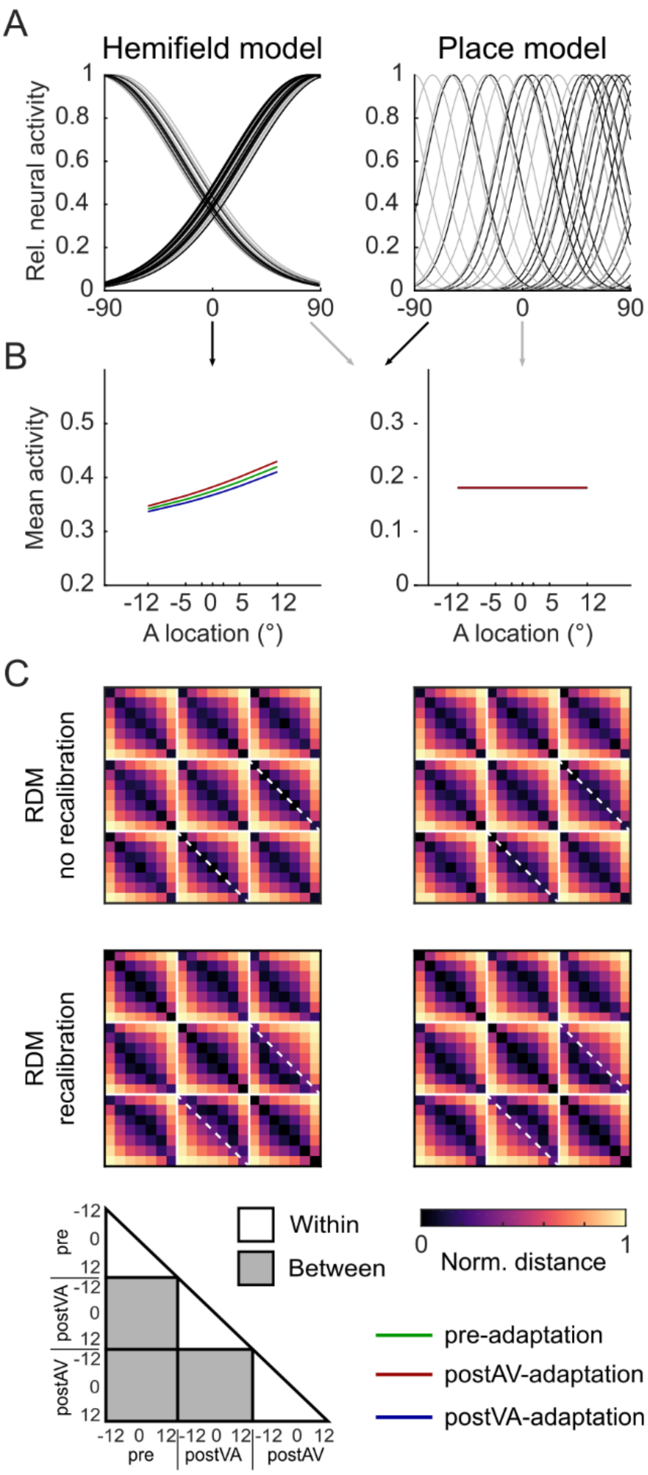

**Figure S1. Comparison of spatial models based on hemifield and place codes.**

**(A)** Neural models: The hemifield model encodes spatial location in the relative activity of two subpopulations of neurons each broadly tuned either to the ipsi- or contra-lateral hemifield. The ratio of the ipsi- and contralaterally tuned neurons was set to 30%/70% (dark grey) and 50%/50% (light grey). The place code model encodes spatial location in the activity of a large population of neurons each narrowly tuned to a particular spatial location. We sampled the means of these tuning functions across neurons either uniformly (light grey) or non-uniformly, i.e., with a contralateral bias (dark grey) over space. **(B)** Predicted mean BOLD-response as a function of sound location along the azimuth in a left hemisphere region for pre- (green), postVA- (blue) and postAV-adaptation (red). Both the spatial hemifield model and the place code model predicts a BOLD-response that increases across spatial locations only when the ratio across the two populations (hemifield) or the means of the narrow tuning functions (place code) are biased towards the contralateral hemifield. **(C)** Predicted representational dissimilarity matrices (RDM) based on the individual model neural activity profiles across spatial locations (-12° to 12°) and experimental phases (pre-, postVA-, and postAV-adaptation). We simulated RDMs from the spatial hemifield (left) and the place code (right) model for (i) top row: no recalibration, i.e., without a representational shift and (ii) bottom row: with recalibration, i.e., with a representational shift. Both models make nearly indistinguishable predictions for the RDMs regardless of exact parameter settings (e.g., uniform vs. non uniform sampling). Solid white lines delineate the sub-RDM matrices that show the representational dissimilarities for different stimulus locations within and between different experimental phases. Diagonal dashed white lines highlight the RDM dissimilarity values for two identical physical locations of the postAV- and the postVA-adaptation phases. Comparing the RDMs with and without recalibration along those dashed white lines shows how the shift in spatial representations towards the previously presented visual stimulus alters the representational dissimilarity of corresponding stimulus locations in postAV- and postVA-adaptation phases, while the off-diagonals show the dissimilarity values for neighbouring spatial locations.

#### References

1. Quax, S. C., Dijkstra, N., van Staveren, M. J., Bosch, S. E. & van Gerven, M. A. J. Eye movements explain decodability during perception and cued attention in MEG. *NeuroImage* **195**, 444–453 (2019).
2. Smith, P. L. & Little, D. R. Small is beautiful: In defense of the small-N design. *Psychon. Bull. Rev.* 1–19 (2018) doi:10.3758/s13423-018-1451-8.
3. Rohe, T. & Noppeney, U. Reliability-Weighted Integration of Audiovisual Signals Can Be Modulated by Top-down Attention. *eNeuro* **5**, ENEURO.0315-17.2018 (2018).
4. Nili, H. *et al.* A toolbox for representational similarity analysis. *PLOS Comput. Biol.* **10**, e1003553 (2014).
5. Werner-Reiss, U. & Groh, J. M. A rate code for sound azimuth in monkey auditory cortex: implications for human neuroimaging studies. *J. Neurosci.* **28**, 3747–3758 (2008).
6. Wood, K. C., Town, S. M. & Bizley, J. K. Neurons in primary auditory cortex represent sound source location in a cue-invariant manner. *Nat. Commun.* **10**, 1–15 (2019).
7. Schlack, A., Sterbing-D'Angelo, S. J., Hartung, K., Hoffmann, K.-P. & Bremmer, F. Multisensory Space Representations in the Macaque Ventral Intraparietal Area. *J. Neurosci.* **25**, 4616–4625 (2005).
8. Salminen, N. H., May, P. J. C., Alku, P. & Tiitinen, H. A population rate code of auditory space in the human cortex. *PLOS ONE* **4**, e7600 (2009).

9. Ortiz-Rios, M. *et al.* Widespread and Opponent fMRI Signals Represent Sound Location in Macaque Auditory Cortex. *Neuron* **93**, 971–983 (2017).
10. Diedrichsen, J., Yokoi, A. & Arbuckle, S. A. Pattern component modeling: A flexible approach for understanding the representational structure of brain activity patterns. *NeuroImage* **180**, 119–133 (2018).
11. Jeffreys, H. *Theory of probability*. (Oxford University Press, 1961).
